# Modeling of protein conformational changes with Rosetta guided by limited experimental data

**DOI:** 10.1101/2022.02.14.480383

**Authors:** Davide Sala, Diego del Alamo, Hassane S. Mchaourab, Jens Meiler

## Abstract

Conformational changes are an essential component of functional cycles of many proteins such as transporters or receptors. Despite their importance, the characterization of functionally-relevant structural intermediates is often a hard task for both experimental and computational methods. This limitation can be at least partially relieved by integrating experimental data within computational approaches. Such integrative structural biology strategies can yield an ensemble of models consistent with experimental data. While the macromolecular modeling suite Rosetta has been extensively used to predict protein structures *de novo* using various types of experimental data, an all-purpose conformational change modeling approach has been absent from this suite. Here, we introduce and benchmark ConfChangeMover (CCM), a new Rosetta Mover tailored to model conformational changes in proteins using sparse experimental data. CCM is able to rotate and translate secondary structural elements and modify their backbone dihedral angles in regions of interest. We benchmarked CCM on soluble and membrane proteins with sparse experimental double electron-electron resonance (DEER) restraints and simulated Cα - Cα distance restraints, respectively. In both benchmarks, CCM outperformed state-of-the-art Rosetta methods, showing that it can model a diverse array of conformational changes. In addition, the Rosetta framework allows a wide variety of experimental data to be integrated with CCM, thus extending its capability beyond DEER restraints. Thus, our method will contribute to the biophysical characterization of protein dynamics.

## INTRODUCTION

Proteins exert a variety of functions that lead to fine-tuned control of biological processes. These functions rely on conformational changes of varying degrees, implying that a single static structure is insufficient to describe a molecular mechanism. While some proteins, such as certain enzymes, might require only small conformational changes for function, other proteins, and in particular membrane proteins such as transporters or receptors, need the motion of a number of structural elements to drive function. Despite great advances in experimental methods for protein structure determination [1–3], the complete characterization of multiple conformational states remains a challenge. Because membrane proteins function through shifts in ensembles of conformations, X-ray crystallography or electron microscopy provide only snapshots of selected conformations. Spectroscopic techniques such as Nuclear magnetic resonance (NMR) or electron paramagnetic resonance (EPR) are better suited to monitor shifts in ensembles of states but typically yield limited data that incompletely define the ensembles. Computational methods are challenged by the large space of conformations that are similar in energy and thus difficult to distinguish with imperfect energy functions. Although advances have recently been made in the prediction of proteins structure from sequence [4–6], including those in multimeric complexes [7–10], the prediction of conformational ensembles and protein dynamics remains a fundamental challenge [11].

Computational techniques such as molecular dynamics (MD) simulations have been extensively used to dissect mechanistically important aspects of protein motion [12]. Recent advances made it possible to reach large spatial and time scales that, in combination with atomic details, can provide insight into mechanisms of conformational changes [13]. Nevertheless, large conformational transitions often occur at time scales beyond those achievable with MD, and consequently experimental measurements must be used to collect information on populations at equilibrium [14]. Depending on the technique, these observables may represent Boltzmann-weighted averages of multiple sampled conformational states and may be limited by poor spatial/time representation [15,16]. In both cases, the obtained macroscopic measurements have the potential to unveil structural transitions between distinct conformations within the functional cycle.

Markov Chain Monte Carlo (MCMC) molecular modeling approaches can complement experimental techniques by determining meaningful three-dimensional structures consistent with limited experimental data [17,18]. Integrative structural biology combines structural information coming from multiple experimental sources, thus compensating for technique-specific shortcomings that result in data that is noisy, ambiguous, and/or unevenly distributed. Several molecular modeling programs have been developed that incorporate purpose-tailored modeling protocols in which experimental information is used as restraints to drive conformational sampling [19–22]. Among them, the macromolecular modeling suite Rosetta has been widely adopted for this purpose, and a variety of applications have been developed addressing diverse modeling tasks [23]. In a typical Rosetta pipeline, input poses alternate between stochastic modification using Movers and evaluation using one of several scoring functions. In contrast with MD, which relies on the application of Newtonian forces and is thus limited by the duration of femtosecond-level time steps, MCMC approaches rely on probabilistic sampling to collect models and have, in the case of Rosetta, no inherent timescale [24]. Tailored sampling of input poses using MCMC can, as a result, overcome high-energy barriers in the energy landscape that might otherwise prevent physiologically relevant conformers from being sampled. For example, in proteins that use rigid-body movements to alternate between multiple states, such as enzymes, receptors and transporters, applying rigid-body rotations and translations to secondary structural elements (SSEs) may allow alternative conformations of interest to be modeled. However, such motions are generally achieved by the *ad hoc* application of one of several other modeling methods, such as methods used for homology modeling or *de novo* structure prediction; a unified approach for conformational change modeling of proteins with diverse topologies using experimental data is still missing.

Here, we introduce ConfChangeMover (CCM), a new Rosetta Mover specifically developed to model conformational changes in proteins using limited experimental data. By employing the same ”broken-chain kinematics” strategy previously used for high-accuracy homology modeling [25], CCM is able to rotate and translate SSEs and modify their backbone dihedral angles in regions of interest without affecting the rest of the structure [26]. We discuss several novel strategies designed to maximize structural similarity to the starting conformation in torsion space while exploring a diverse array of conformations in Cartesian space. CCM was benchmarked in two stages. First, conformational changes were evaluated in a panel of soluble proteins with multiple experimental structures using simulated Cα - Cα distance restraints. Then, experimental distance restraints collected using EPR double electron-electron resonance (DEER) were used to demonstrate how CCM can be applied to modeling of conformational change in large membrane proteins such as receptors and transporters. In both benchmarks, CCM outperformed state-of-the-art Rosetta methods to model conformational changes driven by sparse restraints, suggesting that it can be used to model proteins with a wide variety of topologies.

## RESULTS

We introduce first the two benchmark sets used in this study. One benchmark was carried out on soluble proteins using simulated distance restraints between alpha carbons, whereas the second benchmark was carried out on membrane proteins using experimental EPR restraints. Next, we detail the ConfChangeMover (CCM) protocol. This is followed by a discussion of the accuracy of the models generated by CCM, which are compared to those generated using the Rosetta backbone fragment insertion protocol. Finally, we conclude with results of a membrane protein benchmark using experimental restraints, and discuss the advantages and limitations implied by these results.

### Benchmark on soluble proteins

Seven topologically dissimilar proteins were selected from previous benchmarks on modeling conformational changes and used to determine the effectiveness of CCM [27,28] (Table 1). We first modeled all missing residues using RosettaCM [26]. Distance restraints between Cα atoms were chosen using a previously published restraint-picking algorithm and derived from the target structure [28,29]. The Rosetta scoring functions *score3* and *score4_smooth_cart* were used for stages 1 and 2 of CCM, respectively. We compared CCM to SingleFragmentMover (SFM), which introduces fragment insertion into the backbone; 50,000 rounds of 3-mer fragments insertions were performed with GenericMonteCarlo mover and *score3* scoring function. Fragments were collected using the Robetta web server as previously described [30]. One hundred models were generated for each of the 14 conformational transitions in the benchmark. Cα root mean squared deviation (RMSD) calculations were limited to each target protein’s mobile regions.

**Table 1.** Soluble proteins modeled with simulated C_α_-C_α_ distance restraints.

| Protein (*Organism*) | Conformations (PDB ID) | No of Restraints | No of Modeled Residues | References |
| --- | --- | --- | --- | --- |
| Adenosylcobinamide kinase (*Salmonella enterica*) | Apo (1CBU) - Holo (1C9K) | 18 | 33 | [31,32] |
| Pol alpha DNA polymerase (*Escherichia phage RB6*) | Holo (1IG9) - Apo (1IH7) | 30 | 111 | [33] |
| Glutamine-binding protein (*Escherichia coli*) | Holo (1WDN) - Apo (1GGG) | 21 | 248 (full length) | [34,35] |
| Leucine-binding protein (*Escherichia coli*) | Holo (1USI) - Apo (1USG) | 22 | 121 | [36] |
| DNA polymerase I (*Thermophilus aquaticus*) | Open (2KTQ) - Active (3KTQ) | 26 | 69 | [37] |
| Aspartate aminotransferase (*Gallus gallus*) | Closed (1AMA) - Open (9AAT) | 28 | 167 | [38,39] |
| Lactoferrin (*Homo sapiens*) | Holo (1LFG) <-> Apo (1LFH) | 21 | 160 | [40,41] |

### Benchmark on membrane proteins

For this purpose, we used data collected using double electron-electron resonance (DEER) EPR spectroscopy, which measures distance distributions between nitroxide spin labels attached to the protein backbone. Proteins in the benchmark set were entirely alpha-helical in the transmembrane domain and include the G-protein coupled receptor (GPCR) Rhodopsin and the transporters LeuT [42–44], Mhp1 [45,46], and vSGLT [47–49] (Table 2). We note that unlike the soluble protein benchmark described above, we only sampled conformational changes when sufficient experimental data were available, which excluded both the outward-to-inward transition in LeuT (due to the small number of experimental measurements collected in the presence of the background mutations required to stabilize the target conformation) and the inward-to-outward transition of vSGLT (due to the lack of an experimental outward-facing structure). To serve as a starting point when modeling the outward-to-inward transition in vSGLT, we used a previously published outward-facing homology model generated from its homolog SiaT [49].

**Table 2.** Conformational transitions of membrane proteins modeled with experimental EPR DEER distance restraints.

| Protein (*Organism*) | Starting Conformation (PDB ID) | Target Conformation (PDB ID) | No of Restraints | No of helical Residues modeled | References |
| --- | --- | --- | --- | --- | --- |
| Rhodopsin (*Bos taurus*) | Active (2X72) | Inactive (1GZM) | 10 | 96 | [50,51] |
| Rhodopsin | Inactive (1GZM) | Active (2X72) | 10 | 96 |  |
| LeuT (*Aquifex aeolicus*) | Inward-facing (3TT3) | Outward-facing (2A65) | 11 | 167 | [42–44] |
| vSGLT (*Vibrio parahaemolyticus*) | Outward-open<sup>†</sup> (5NV9) | Substrate free (2XQ2) | 11 | 241 | [47–49] |
| Mhp1 (*Microbacterium liquefaciens*) | Outward-facing (2JLN) | Inward-facing (2X79) | 14 | 209 | [45,46] |
| Mhp1 | Inward-facing (2X79) | Outward-facing (2JLN) | 14 | 201 |  |

<sup>†</sup> The vSGLT OF conformation was generated from the X-ray structure of the homolog SiaT.

Simulated DEER distance restraints were computed with the MDDS method implemented in the charmm-gui webserver [52–55].The resulting simulated DEER distances as well as the simulated Cα – Cα distance restraints were applied only on the residue pairs of experimental restraints. Experimental DEER distances of LeuT-fold transporter proteins have previously been collected using EPR [44,49,56,57]. DEER restraints were provided to Rosetta through the RosettaDEER module either as quadratic functions centered on the median values of each distance distribution or as flat-bottom potentials centered on the median values, plus or minus 5 Å, of each distance distribution [58]. The regions of the proteins spanning the membrane were predicted with OCTOPUS [59]. Rigid-body segments were modified to include all the residues involved in restraints. Fragments were collected as described above for soluble proteins. For CCM, we set stages 1 and 2 to consist of 30,000 rounds and 4,000 rounds, respectively. For SMF, 30,000 rounds of 3-mer fragments insertions were performed with GenericMonteCarlo mover. For all the runs, the Rosetta scoring function *stage2_membrane* was used [60]. For RosettaCM, *stage2_membrane* was used in stage1 and stage2, whereas *stage3_rlx_membrane* was used in stage3.

### ConfChangeMover Workflow

The ConfChangeMover (CCM) was developed in Rosetta and executed in RosettaScripts [61]. It samples candidate structural models using a two-stage strategy mainly consisting of rigid-body movements of SSEs followed by loop closure. Prior to the first stage, the input structure is converted to a coarse-grained model with side chains replaced by centroid pseudo-atoms. Cutpoints are introduced at residues located on loops connecting pairs of rigid bodies, or SSEs, which consist of either α-helices or β-sheets identified using the Dictionary of Secondary Structure of Proteins (DSSP) [62]. The introduction of cutpoints at connecting loop regions prevents local perturbations from propagating to the rest of the protein via the “lever-arm effect” [63].

To account for the relative invariance of protein dihedral angles during conformational change modeling, circular sigmoidal restraints are added to the model’s *ϕ* and *ψ* angles based on either the starting conformation or a separate model provided by the user (Eq 1).

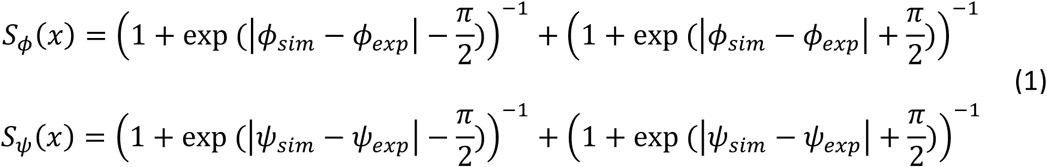

Here *ϕ*_*sim*_/*ψ*_*sim*_ are the *ϕ* and *ψ* angles observed in the model and starting conformation, respectively. These restraints penalize changes in backbone dihedral angles up to, but not beyond, a certain rotation angle. The use of sigmoid functions generally engenders restraint violations that are limited to relatively few degrees of freedom; in contrast, harmonic restraints instead typically distribute restraint violations throughout the features. Thus, these functions quantify the expectation that most, but not all, dihedral angles remain unchanged during isomerization; similar restraints have previously been applied when using ambiguous data such as residue coevolutionary couplings [64]. These restraints are added to residues belonging to SSEs in stage 1 and to loop regions in stage 2. The weight of the *dihedral_constraint* score term, which balances the contribution of these restraints relative to the other components in the scoring function, was set to 1.0 and 0.1 during the first and second stages, respectively.

#### Stage 1: Rigid-body structural perturbation

During the first stage, several types of perturbations are randomly introduced. First, SSEs can be moved in isolation, with rotation angles and translation vectors randomly drawn from normal distributions with user-defined standard deviations. In the soluble protein benchmark discussed below, these comprised 32% of all moves sampled in stage 1, and the standard deviation of rigid-body rotations and translations were 10° and 1.0 Å, respectively. Second, up to *N* − 1 SSEs close in space can be moved in tandem, where N is the number of SSEs in the model (accounting for 50% of moves). Third, helices may be twisted along their axis by a randomly chosen angle drawn from a normal distribution with a user-defined standard deviation (8% of moves). Finally, dihedral angles of three-residue stretches can be modified to match those of randomly chosen sequence fragments obtained from the PDB (10% of moves). In the soluble protein benchmark, stage 1 consisted of 50,000 total moves.

Throughout stage 1, loops are allowed to break and move independently at pre-specified cutpoints as described above. Nevertheless, plausible topologies are implicitly enforced throughout stage 1 with distance constraints between SSEs adjacent in sequence that prevent them from being separated by distances that cannot be bridged by the loops between them. The maximum allowable distance between the C-terminus of one SSE and the N-terminus of next SSE was set to (2.65\**n*res + 2.11) Å where *n*res is the number of residues in the loop, which has previously been used in *de novo* protein folding with implicit loops [65].

#### Stage 2: Loop closure and structural minimization

Between the first and second stages, coordinate constraints are applied to the Cα atoms of all residues in SSEs, thus preventing large-amplitude distance changes from being introduced during loop closure. During the second stage, loops are closed using an approach designed for multi-template homology modeling [66]. Briefly, nine-residue sequence fragments obtained from the PDB are superimposed in Cartesian space on regions of the protein with chain breaks. During the final 25% of Stage 2, this superimposition is followed by Cartesian minimization using the L-BFGS minimization algorithm for five iterations, which further reduces the size of the gaps [67,68]. To increase model diversity, these superimpositions are periodically applied to regions of the protein that fail to have gaps (50% of moves in our soluble protein benchmark). Additionally, to maintain similarity to the starting structure, some of these nine-residue fragments are replaced by nine-to-fifteen residue fragments derived from the starting conformation (80% of moves in our soluble protein benchmark). In conjunction with the sigmoidal potentials described above, these moves minimize the magnitude of the changes undertaken by loop regions, which were otherwise observed to undergo dramatic movements inconsistent with experimental observations. We found that 5,000 moves were sufficient to close all loops. Finally, at the end of stage 2, the entire model is minimized using 2,000 iterations of the L-BFGS algorithm.

### Modeling conformational changes of soluble proteins with simulated distance restraints

We first benchmarked ConfChangeMover (CCM) using simulated harmonic distance restraints on a dataset of seven soluble proteins with structures that have been experimentally determined in two or more conformations (Table 1). Proteins in the benchmark set were selected on the basis of their modes of conformational isomerization, wherein loops and secondary structure elements (SSEs) undergo structural changes that are unlikely to be sampled by rotation of backbone dihedral angles. CCM was compared to SingleFragmentMover (SFM) [69,70], a Monte Carlo-based approach that samples backbone torsions by fragment insertion. Additionally, CCM was further run without restraints to ascertain the effectiveness of the simulated distance restraints. To evaluate modeling accuracy, we calculated each model’s root mean squared deviation (RMSD) across Cα atoms relative to the target conformation. Overall, one hundred models were generated using each approach across each of 14 conformational transitions, with sampling and RMSD calculations focused on mobile regions of the protein structure.

The benchmark included conformational changes of varying degrees and complexity that can be visualized by superimposing the two native conformations of each protein (Figure 1). The two conformations of Adenosylcobinamide kinase differ mostly in a small helix position that can be recapitulated by a rigid-body movement and subsequent loops remodeling, whereas most of the Pol alpha DNA polymerase transition involves a rigid-body movement of two long helices and a change in their curvature. More complex movements are observed in Glutamine-binding protein that cover the whole structure. In Leucine-binding protein, the most relevant movement involves a helix switch between attached and detached state requiring a translation of several angstroms and a rotation of more than 180°. The DNA polymerase I transition includes only a small portion of the protein that includes significant dihedral changes, leading the open state to be more unfolded than the active state. The Aspartate aminotransferase protein predominantly undergoes movement in a single helix that experiences a substantial translation and bending. Lastly, Lactoferrin is characterized by a rigid-body movement of a whole domain that is discontinuous in sequence space.

**Figure 1.**
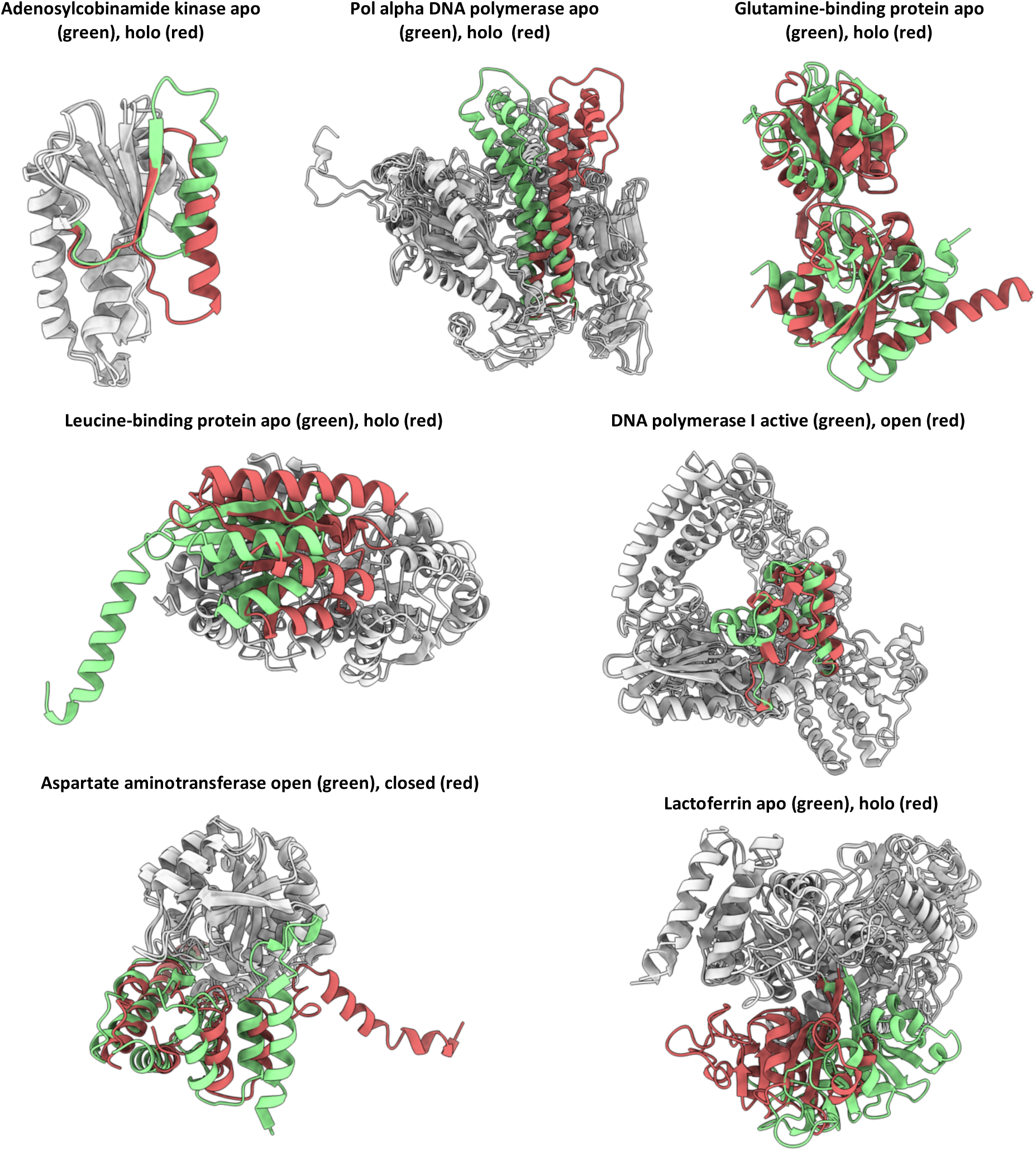
Conformational changes of soluble proteins in the benchmark. Regions modeled are in red or green. Regions not modeled are in gray.

The accuracy in modeling conformational changes of folded regions was assessed by calculating Cα RMSD of mobile residues forming helices or β-sheets (Figure 2). In the absence of restraints, the majority of models sampled using CCM were close to the starting conformation (dashed line in the plot) suggesting that starting structures generally occupied an energy minimum that may be difficult to escape. The inclusion of distance restraints generally led to models with lower RMSD values. By contrast, modeling conformational transitions using SFM with restraints led to models with a wide range of RMSD values; in many cases, these RMSD values increased compared to that of the starting conformation. Overall, CCM outperformed SFM in all the transitions with one exception, the modeling of DNA polymerase I from active to the more unfolded open conformation where SFM generated a slightly better ensemble than CCM. Even in this case, however, SFM generated far more outliers with high RMSD values than CCM.

**Figure 2.**
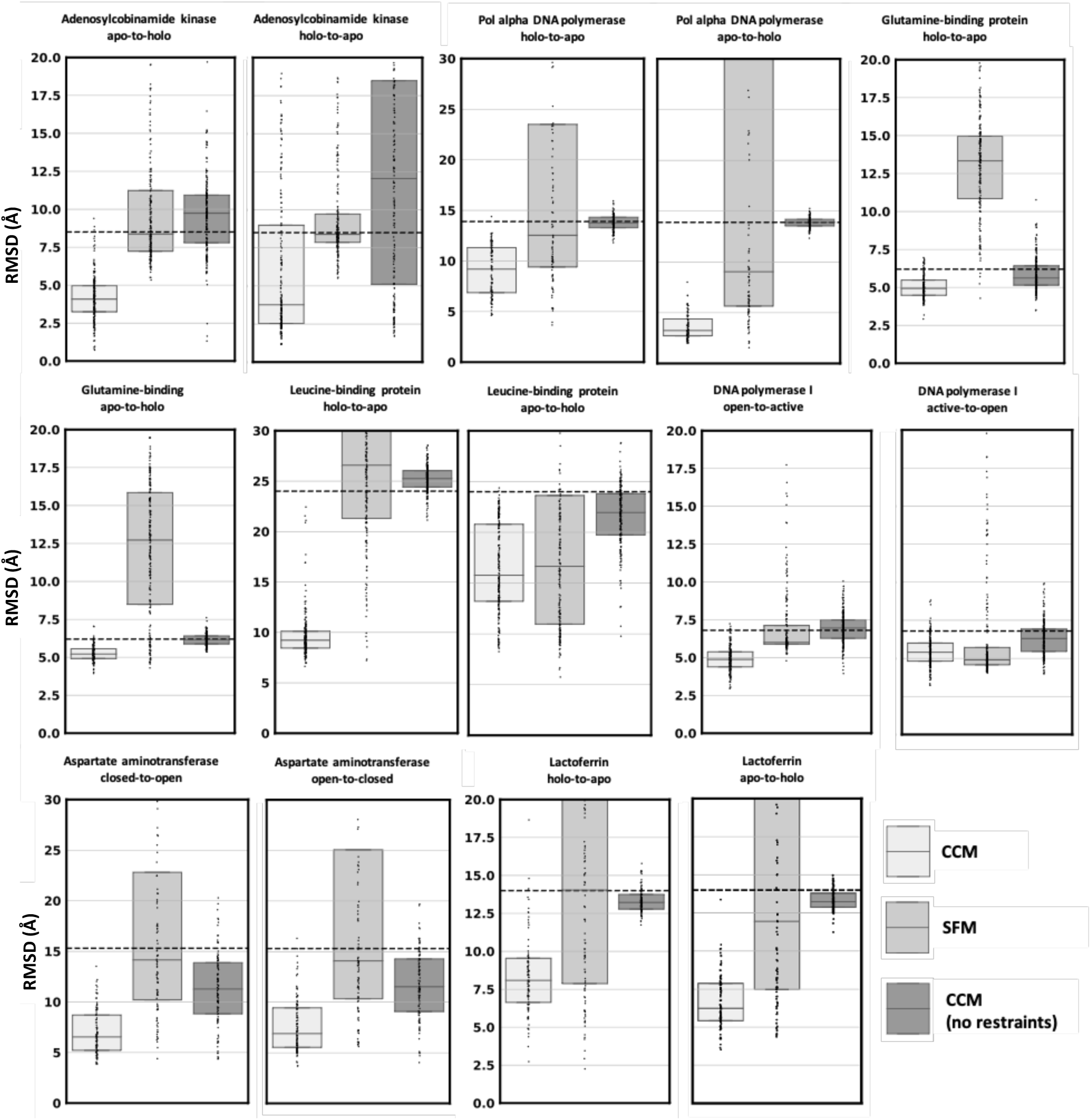
CCM performance in modeling conformational changes of SSEs using simulated Cα-Cα distance restraints. Dots represent the real distribution of RMSD values. The RMSD between the two native conformations is shown as a dashed line.

Including loop regions in Cα RMSD measurements generally led to increases in median RMSD values (Figure S1), which was unsurprising given that loops are among the most conformationally flexible regions in protein structures. In several cases, their inclusion during RMSD calculations yielded RMSD values that more closely approximated those of the starting models. For example, when modeling the apo-to-holo transition of Glutamine-binding protein, RMSD values improved on average by 0.9 Å when limiting RMSD calculations to residues on helices or sheets, but only 0.1 Å when loop regions were included in the calculation. Similarly, the RMSD values of models recapitulating the closed-to-open transition in DNA polymerase I were markedly similar to that of the starting structure only when loop regions were considered. We therefore suspected that CCM was generally more accurate in modeling SSEs rather than unfolded regions. To investigate this hypothesis, we computed the RMSD change of SSEs and loops with respect to the starting value observed between native conformations (Figure S2). Both CCM and SFM increased RMSD of models when loops are included in the measurements. However, models generated by CCM had a RMSD on average 1.2Å higher than SSEs only, a value that is double that of models generated by SFM. These results indicate that CCM was less effective at modeling disordered regions than at modeling structured regions. An illustrative example of this fact is the small DNA polymerase helix (residues range 638-647), which alternates between almost completely unfolded in the open conformation to fully folded in the active state (Figure S3). We found that CCM more accurately modeled the unfolded-to-folded transition than the folded-to-unfolded, whereas SFM left that region mostly unchanged in both cases.

Visual inspection of the models generated using CCM with the lowest RMSD values suggested that this method could more accurately model rigid-body rotations and translations of α-helices than either helical bending or manipulation of β-sheets (Figure 3). For example, models of the glutamine- and leucine-binding proteins, which have α/β topologies, were both more accurate on helices than on β-sheets. Similarly, whereas models of Adenosylcobinamide superimpose well with their target structures, their β**-**strands appear totally or partially unfolded in the apo and holo model, respectively. Similar observations were made in models of Lactoferrin, which may be due to the omission of full-atom refinement. The Pol alpha DNA polymerase holo conformation was modeled with an outstanding accuracy of 2 Å RMSD, whereas the apo conformation characterized by two long curved helices was less accurate. In agreement with previous considerations, DNA polymerase I models accurately recapitulated conserved SSEs but missed the dihedral changes needed to sample the small helix switching folding state.

**Figure 3.**
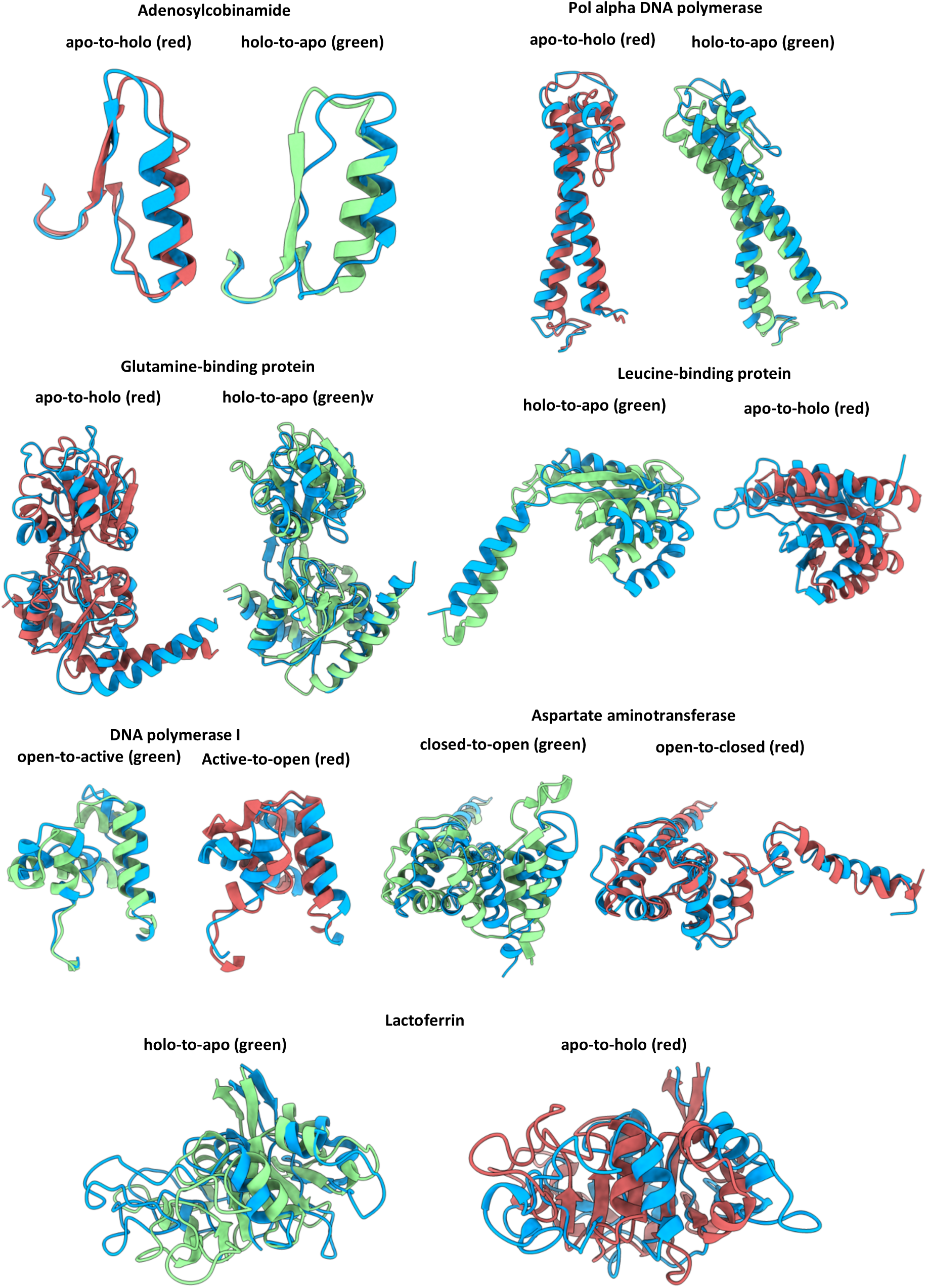
Superimposition between the best model generated by CCM with simulated distance restraints and the corresponding target structure. The target native conformation is in green or red, following the color code used in Fig 1. Models are in blue.

In summary, the data suggest that CCM is a robust and modular sampling method capable of accurately modeling conformational changes guided by simulated distance restraints. Its conservative sampling approach, which prioritizes rigid-body rotations and translations of SSEs, allows it to easily outperform SFM, which has previously been used to sample conformational changes in Rosetta [66]. The primary drawback we observed was the modeling of new loops conformations, which was difficult mainly due to stage 2 parameters that erred toward conservative sampling. It should be noted that only 100 models were generated for each transition, and no attempt was made to fine-tune the sampler for each test case (see *Discussion* for modifications that can be introduced by the user). Indeed, protein-specific modifications to the protocol may ameliorate the slightly worse performance observed when modeling structural changes such as helical bending or unfolding events. We surmize this is likely due to the limited extent to which sequence fragments are used when modifying backbone dihedral angles. Similarly, more accurate loop conformations can be generated by simply using less conservative stage 2 parameters (e.g. decreasing the contribution of the appropriate score terms) while increasing the number of stage 2 models. Thus, while Cα RMSD values approached 2-6 Å in some cases of our benchmark, performance could almost certainly be improved across the board by either modifying the sampling parameters, increasing the duration or aggressiveness of sampling, or following CCM with full-atom minimization.

### Modeling conformational changes of membrane proteins with experimental EPR DEER distance restraints

Having assessed the capacity of CCM on soluble proteins with simulated distance restraints, we evaluated its effectiveness at modeling membrane proteins using previously published experimental data (Table 2). The benchmarked proteins undergo divergent modes of conformational change to facilitate transmission of signals or translocation of substrates across the membrane (Figure 4). As with the soluble protein benchmark, we only allowed specific regions to move where experimental data are available. The only exception was Rhodopsin, for which transmembrane (TM) helices 5, TM6 and TM7 are the three helices most involved in mediating protein activation and deactivation. In Mhp1, interconversion between the IF and OF states was mediated by rigid-body movement of the hash domain (TM3, TM4, TM8, and TM9), as well as bending of TM5. For LeuT, helices in the bundle domain (TM1, TM2, TM6, and TM7), as well as TM5 and the amphipathic helix on the loop connecting TM7 and TM8, were permitted to move, while the positions of the hash domain and TM10-12 were fixed. For vSGLT, the conformational change suggested a mechanism of alternating access involving helices throughout the entire protein. Among them, TM11 undergoes an extensive change that results in a highly bent helix in the target substrate-free (SF) conformation.

**Figure 4.**
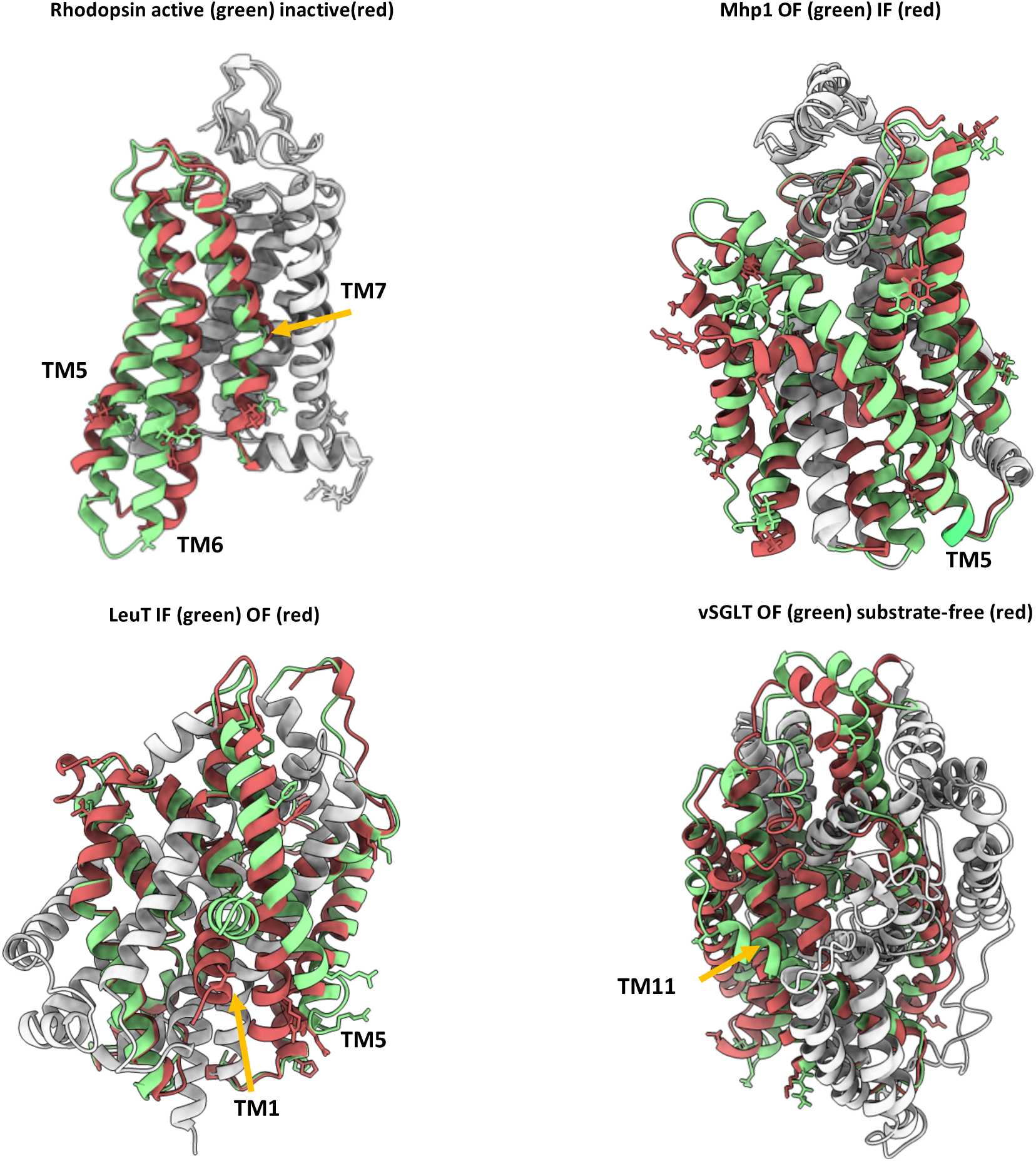
Benchmarked conformational changes of membrane proteins. Regions modeled are shown in red or green. Regions not modeled are in gray. Restrained residues are shown as sticks.

Experimental distance data was incorporated as restraints using the RosettaDEER module [58]. RosettaDEER models the ensemble of nitroxide spin probe conformations using coarse-grained depictions (called pseudorotamers) designed to maximize computational efficiency. After removing pseudorotamers that clash with the protein, the average distance of the sampled distribution between pairs of ensembles was calculated and compared to the experimental average distance using two different scoring approaches. First, a direct comparison was carried out, and scores reflected the squared deviation between the sampled and experimental average distance values. Second, only deviations beyond 2.5 Å were penalized, permitting models to adopt conformations that may not be in full agreement with the data without penalty. Additionally, to ascertain the effect of these experimental restraints, we provided the median distance value simulated using the method MDDS [52,53] that generates distances with a strong correlation with those simulated using RosettaDEER [58]. Finally, to directly compare the contribution of these probe-based measurements to Cα distance restraints, we provided simulated Cα distance restraints between the same residues used for experimental measurements.

In this benchmark, CCM was compared to both SFM and the comparative modeling method RosettaCM [26], which samples conformational movements in a superficially similar way and is also used to model conformational changes and refine protein structures [71,72]. Each of these two methods exclusively used DEER restraints as experimental average distance values.

All approaches generated 1000 models, which were then compared to the target structures by RMSD (Figure 5). As with the soluble benchmark set discussed above, these data illustrate how CCM outperforms SFM in modeling each conformational change of interest. SFM generated models with a wide range of RMSD values, and visual inspection of these models indicated partial unfolding. By contrast, RosettaCM appeared to sample conformations that were structurally similar to the starting structure, which is likely due to its emphasis on sampling changes in torsion space and its inability to introduce the rigid-body rotations and translations of interest. In contrast, models generated using CCM showed a broader distribution of RMSD values, indicating that the method sampled SSEs more aggressively. As a result, in most cases this approach generated models with the lowest RMSD values among all methods considered. The two noteworthy exceptions are the outward-to-inward transition in Mhp1 and the inactive-to-active transition in Rhodopsin. However, distributions of models generated with simulated data suggest that this can be at least partially attributable to the quality of the restraints. Experimental restraints provided as median values or as ranges showed a similar performance with the exception of Rhodopsin transitions for which ranges allowed the sampling of alternative conformations closer to the target.

**Figure 5.**
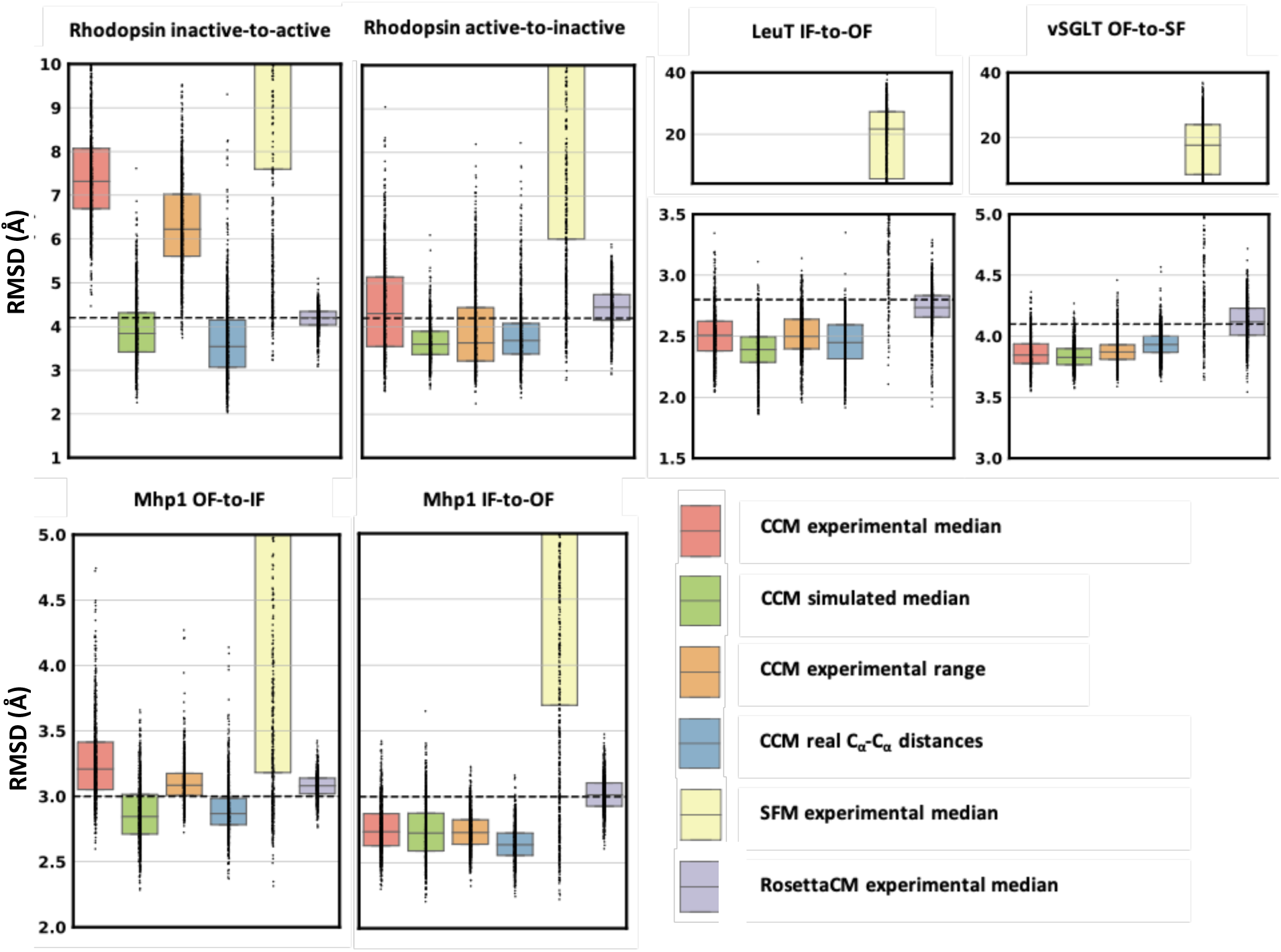
RMSD from target conformation of modeled membrane proteins conformational ensembles. Restraints were provided as median values of the experimental distribution (experimental median) or as a range centered on the median value (experimental range). Dots represent the real distribution of RMSD values. The RMSD between starting and target conformation is shown as a dashed line.

The accuracy of each DEER dataset was evaluated by measuring its average RMSD from the simulated average (Figure S4). Substrate-free vSGLT and OF LeuT conformation featured the lowest and the highest RMSD of 2.7 Å and 4.6 Å, respectively. All the remaining datasets spanned a range between 3.5 Å and 4.0 Å. Active Rhodopsin and IF Mhp1, which were among the most difficult to model with experimental data, showed the broadest distributions of model accuracies. Thus, few highly inaccurate restraints may have been at least partially responsible of poor modeling. To identify the most inaccurate restraints, we computed the difference between experimental and simulated distance for each residue pair restraint (Figure S5). Restraints that stood out in the Rhodopsin active-state dataset included those involving residues 225 (TM5), and residues 241 and 252 (TM6). Analogously, Mhp1 residues 30 and 338 were involved in the most inaccurate restraints of the IF state. The contribution of these residues on the global RMSD of models was then assessed by computing their per-residue RMSD from the target conformation (Figure S6). As expected, all these residues have greater RMSD with respect to the median value of global RMSD of models (dashed line). Most of the Rhodopsin models have also per-residue RMSDs greater than in the starting conformation (red dots).

In addition to experimental restraints, accuracy of models is also affected by the complexity of the conformational change to be sampled. For each transition, we then superimposed the best model sampled with experimental restraints on the corresponding target structure (Figure 6). Not surprisingly, whereas TM6 of the active-state model of Rhodopsin superimposed poorly with the target structure, the Rhodopsin inactive state was modeled with high accuracy. Even the OF-state of Mhp1 was accurately modeled, especially the unbending of TM5, whereas the two small helices carrying the 278-362 restraint superimpose poorly onto the target structure. Further, in modeling the Mhp1 IF-state, the helix carrying residue 30 is significantly misplaced, and the bending of TM5 was not introduced using our approach. In LeuT, which underwent transitions defined by large-amplitude movements in the partially unwound helix TM1 and the fully continuous helix TM5, we found that CCM accurately modeled the former, but not the latter. In particular, the motions of the cytoplasmic end of TM5 were driven by the two poorly restrained residues 185 and 193, leading to partial unfolding in the model with the lowest RMSD. Interestingly, while helical bending was generally difficult to model, the vSGLT model correctly reproduced the challenging TM11 bending, which may be due to steric hindrance surrounding this helix. Thus, in general, the most inaccurate distance distributions in the restraints’ dataset appear to directly reduce the accuracy achieved in modeling the two SSEs containing the two spin-labeled residues.

**Figure 6.**
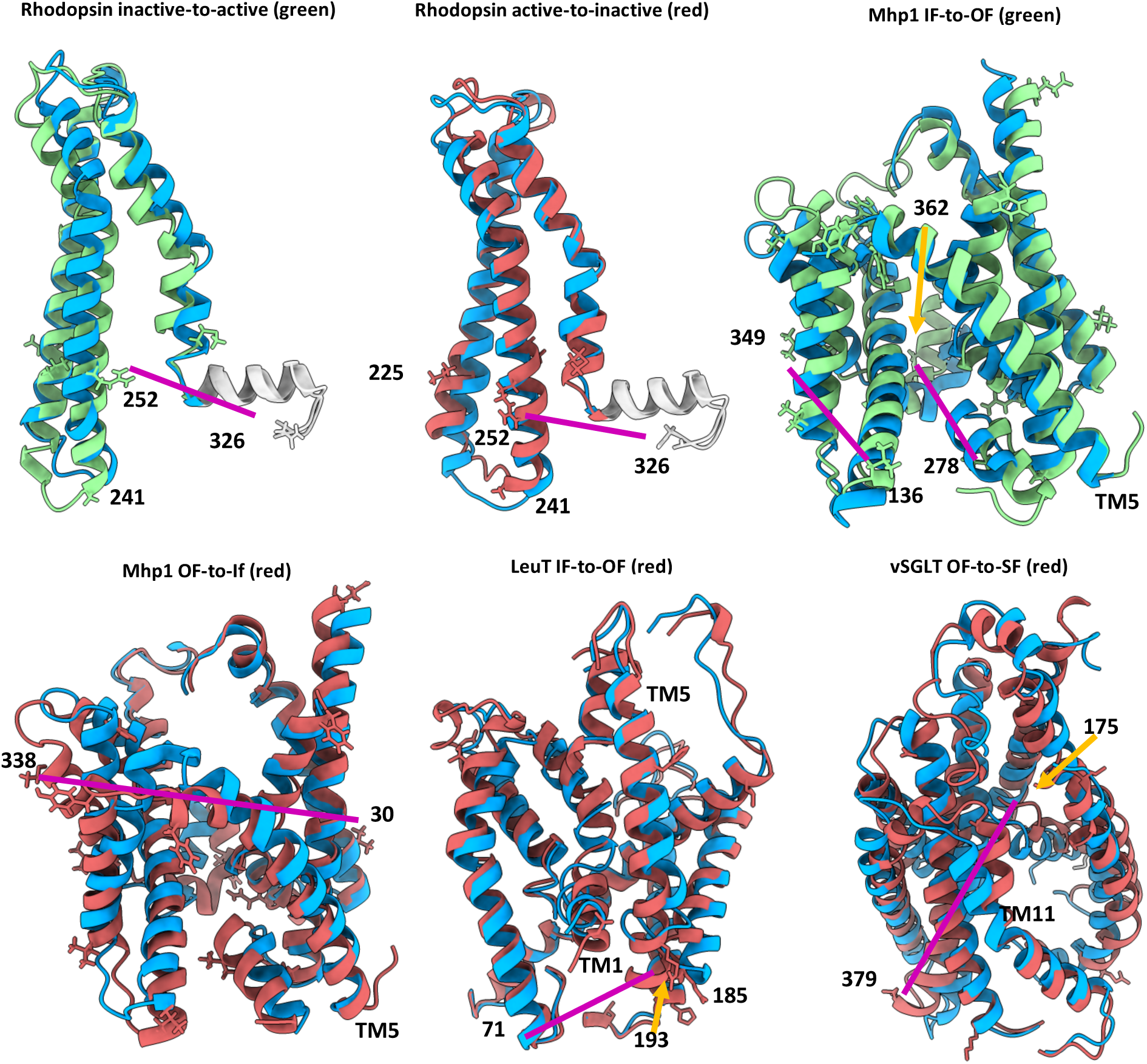
Superimposition between the best model generated by CCM with experimental DEER distances and the corresponding target structure. The target native conformation is in green or red, following the color code used in Fig 4. Rhodopsin helix 8 is in gray. The model is shown in blue. Restrained residues are represented as sticks. Residues involved in restraints featuring a relevant difference between experimental and simulated DEER distances are labeled and mapped on structures.

In summary, CCM outperformed RosettaCM and SFM in modeling conformational changes of membrane proteins driven by high-quality sparse DEER distances. Few highly inaccurate restraints hampered the sampling of accurate models in a couple of conformational transitions that however were successfully modeled with simulated restraints, proving the efficiency of the method. Of note, simulated restraints were applied on the same few residue-pairs of experimental data. As was observed in soluble proteins, conformational transitions involving partial and localized change in the folding state and change in backbone dihedral angles were difficult to sample without extensive fragments insertion. Ultimately, these results suggest that CCM is an effective protocol for conservative sampling of structural models, but that fine-tuning specific to each protein may be necessary to achieve the best results possible.

## DISCUSSION

Here, we described and benchmarked ConfChangeMover (CCM), a new modeling method in the software suite Rosetta that uses experimental restraints to model conformational changes in proteins. The performance of CCM was evaluated in both soluble and membrane proteins using simulated or experimental distance restraints, respectively. In both cases, CCM outperformed existing Rosetta methods that have been previously used to model conformational changes in proteins with a wide variety of topologies [66,73–75], highlighting its versatility and robustness. We believe the main advantage of CCM over other methods stems from its ability to automatically identify, group, and move SSEs as rigid bodies, a task that has been absent from the Rosetta modeling suite. Notably, although CCM allows the user to define sets of SSEs that can only be moved as a group, this option was not used in our benchmark, potentially understating the extent to which this protocol can be used to accurately model conformational changes.

A secondary function of this protocol is fragment insertion to both diversify SSEs in dihedral space and close loops in Cartesian space following rigid-body manipulation of SSEs. Because CCM’s two-stage approach allows loop closure to be decoupled from structural diversification, multiple distinct loop conformations can be quickly generated following the more expensive structural diversification performed during the first stage. Throughout our benchmark, the accuracy achieved in modeling loop regions with simulated distances indicated that the sampling parameters may have been too conservative for accurate modeling. Besides increasing the intensity of sampling, loops can also be modeled afterwards with other Rosetta methods appositely developed for that purpose [76–79], or refined using gradient minimization methods such as FastRelax, which can optimize backbone geometry following fragment insertion [26]. Full-atom refinement could further lead to the correction of small errors, such as the slight unfolding of beta structures observed in some models of soluble proteins.

An advantage of CCM that is not captured by these results is its potential flexibility and modularity. By interfacing with RosettaScripts, CCM provides many options to the user to adjust the protocol in accordance with expectations about the target conformation (see Appendix 1 for more details). In stage 1, the residues in each SSE can be manually defined using residue selectors, which can specify segments and/or change the extension of each segment. In addition, multiple SSEs can be treated as rigid bodies during stage 1. Finally, users can define the number, frequency, and magnitude of each move. During stage 2, the choice of which regions of the protein can be perturbed is also provided and can be expanded to include SSEs which are omitted by default. Additionally, since we found that stage 1 was far more computationally expensive than stage 2, we designed stage 2 to permit multiple output models to be generated from each pose sampled by stage 1. This would have the effect of generating topologically similar models with different loop conformations. However, the results discussed in this report do not use this feature.

It is important to note that despite the parametrization options available to the user, no modifications to the protocol were introduced that accounted for the varying topologies and sizes of protein targets in our benchmark set. Despite this handicap, CCM managed to extensively sample accurate models of soluble proteins using only one simulated distance restraint per twenty residues. Indeed, while the median RMSD improvements often exceeded 5 Å, specialized parametrization would almost certainly lead to further increases in accuracy. Models of membrane proteins generated with simulated restraints had less impressive RMSD improvements; however, the magnitude of their conformational changes were lower than those of soluble proteins, likely precluding comparable improvements in RMSD. We highlight that this benchmark does not account for the placement of experimental restraints in the membrane proteins modeled here, as these were guided by unique scientific questions specific to each protein[80]. By contrast, the soluble protein benchmark used simulated restraints that were chosen using a well-validated algorithm, leading to well-distributed residue pairs covering most of the regions of interest. This methodological difference also explains why conformational changes were modeled in membrane proteins with fewer restraints per residue. Nevertheless, CCM consistently improved the RMSD of any conformational transition modeled, a record that was not shared by the other methods explored here.

While our benchmark was limited to distance restraints, the Rosetta framework allows a wide variety of types of experimental data, potentially from multiple sources, to be used for conformational change modeling tasks. In addition to EPR data, restraints may be obtained from data collected with NMR [73,81], mass spectrometry [82,83], and other sources of geometrical restraints such as FRET and cross-linking. In addition to experimentally derived restraints, computational restrains can also be exploited in modeling conformational changes. Indeed, evolutionary couplings derived from multiple sequence alignments (MSA) can capture protein dynamics by exploring their use as reaction coordinates [84].

In conclusion, CCM facilitates the effective sampling of protein dynamics. This method is well-positioned to deepen our understanding of proteins structural dynamics. Future directions may include structural refinement using residue-residue contacts predicted by machine-learning methods [72] which may facilitate the detection of functional states in proteins spanning multiple heterogeneous conformational states.

## Acknowledgments

We would like to thank Drs. Jeff Abramson and Aviv Paz for providing us with the outward-facing model of vSGLT, Dr. Christian Altenbach for providing the experimental data for Rhodopsin, and Dr. Marion F. Sauer for fruitful discussions. Authors acknowledge funding by the Deutsche Forschungsgemeinschaft (DFG, German Research Foundation) through SFB1423, project number 421152132 and the NIH R01 GM080403, R01 GM129261, R01 HL122010, and R01 DA046138.

## Author contributions

H.S.M., and J.M. conceived the idea. D.S. and D.d.A. designed the framework, wrote the code, performed the calculations and analyzed the data with guidance from H.S.M. and J.M. D.S. and D.d.A. wrote the manuscript and prepared figures, H.S.M. and J.M. further revised the manuscript.

## APPENDIX 1

### Modeling of Adenosylcobinamide kinase apo-to-holo conformational change with distance restraints

In this demo the transition of a small helix and its two adjacent loops (from green to red in the picture) will be modeled with the ConfChangeMover. The small helix in the target structure is longer than in the input conformation and is shifted in the sequence composition, making the modeling quite complex. Its Cα RMSD between native conformations is 10.6 Å. The total Cα RMSD of the region to model (helix + loops) is 9.9 Å. To intensify the sampling of loops, in stage2 three different loops conformations will be created for each helix transition.

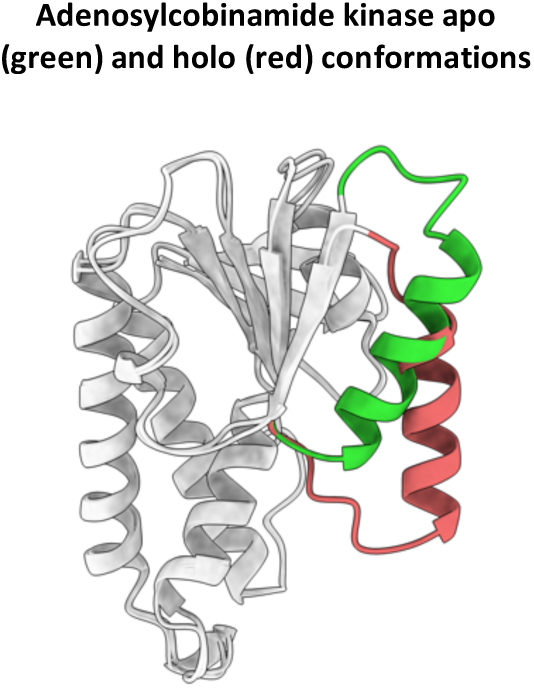

#### Preparation of the input files

The apo (PDB ID 1CBU) and holo (PDB ID 1C9K) structures can be downloaded from ProteinDataBank **REF**. All the residues except range 1-180 of chain A are removed. In the target holo conformation all the missing residues can be modeled by uploading sequence and template in the user-friendly portal https://swissmodel.expasy.org (click on “start modeling” followed by “User template: on the right) **REF**.

Fragments can be collected through http://old.robetta.org **REF**. The website requires the protein fasta sequence to generate 3-mer and 9-mer fragments.

Four Cα distances involving two residues and the termini of the helix have been derived from the target structure to be used as restraints.

All the following input files have been placed inside the “input_files” folder. “1cbu.pdb” and “1c9k.pdb” for input apo and target holo structures, respectively. “aat000_03_05.200_v1_3.txt” and “aat000_09_05.200_v1_3.txt” for 3-mer and 9-mer fragments, respectively. The fasta sequence file “1c9k.fasta”. Restraints are in “1c9k.cst”.

#### RosettaScripts XML file

RosettaScripts XML file used for modeling has been deposited in the “scripts” folder and is shown below. To increase loops variability the frequencies of using template segments and of targeting exclusively gaps have been kept low to 0.1. The “MultiplePoseMover” is required to perform actions on multiple stage2 models out of a single stage1 sampling.

Besides ConfChangeMover, a couple of metrics are calculated: 1) the RMSD from the native structure of the helix (_H suffix) and 2) the RMSD of the whole modeled region (_HL suffix).

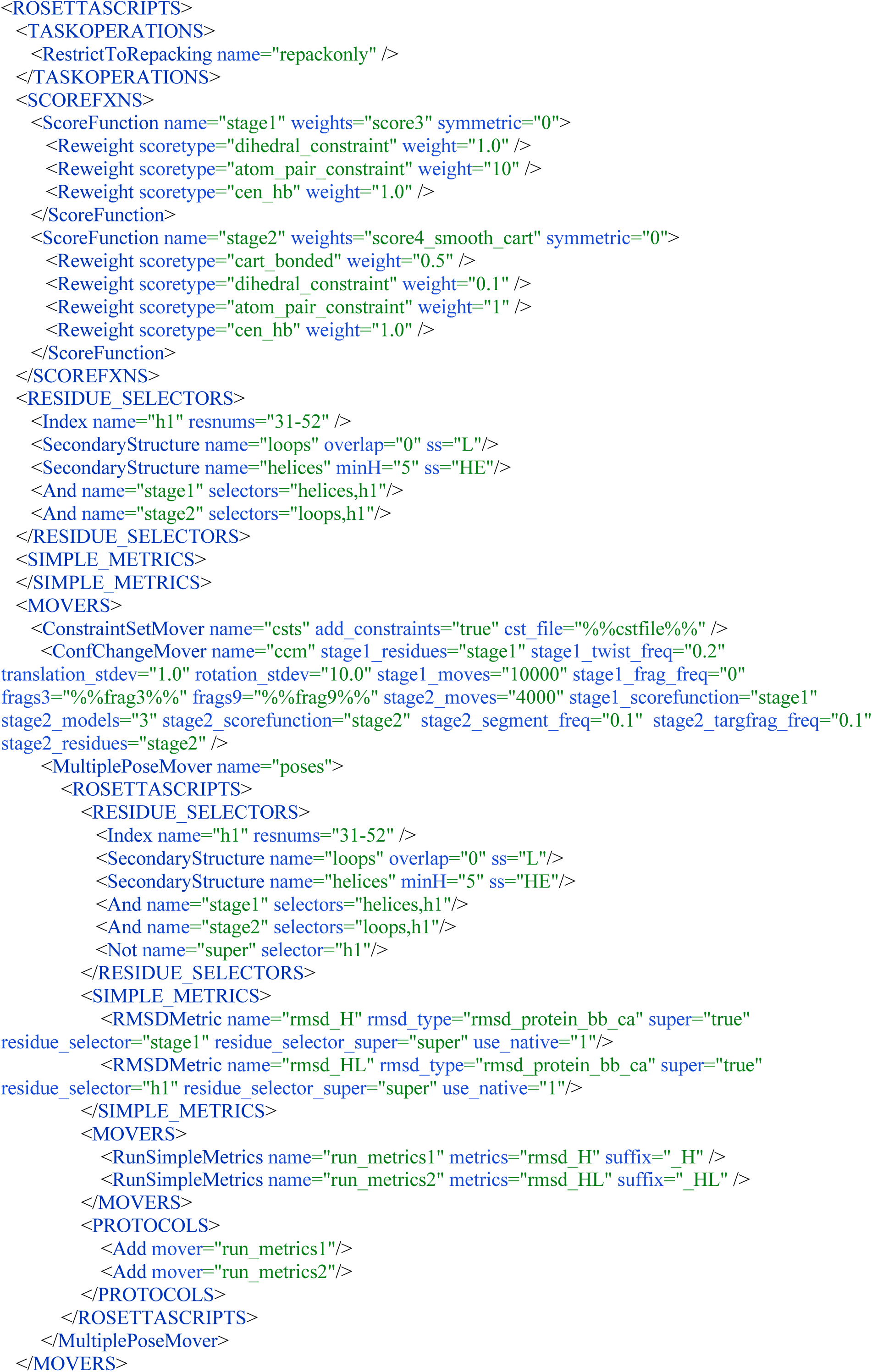
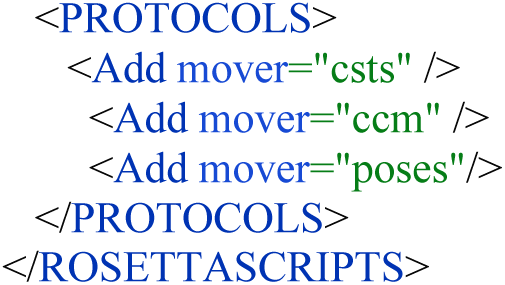

#### Running the command to perform the modeling

By executing the command below from the “working_directory” folder, 9 models are generated in approximatively 20 minutes on a single core. User may need to change the executable depending on the OS used, MAC/linux/Windows. Rosetta executables are located in “path/to/Rosetta/main/source/bin”.

“rosetta_scripts.linuxgccrelease -in:file:s ../input_files/1cbu.pdb -in:file:native ../input_files/1c9k.pdb -parser:script_vars frag3=../input_files/aat000_03_05.200_v1_3.txt frag9=../input_files/aat000_09_05.200_v1_3.txt cstfile=../input_files/1c9k.cst - parser:protocol ../scripts/ccm.xml -out:nstruct 3 -out:path:pdb ./ -mute core.optimization.Minimizer”

#### Analyzing results in output

Depending on the number of models generated user can find the corresponding number of PDB files and a score file inside the “working_directory” folder. The score file contains the RMSD metrics for each model. Pre-generated output files can be found in the “output_files” folder.

### Modeling of Rhodopsin active-to-inactive conformational change with simulated DEER distances

Here, Rhodopsin transition from active to inactive state will be modeled. Such transition involves the three helices TM5-TM6-TM7 that have a Cα RMSD of 4.2 Å, the inclusion of loops does not change RMSD. The cytoplasmic ends of TM5 and TM6 differ in the two protein states. The longer helices in the active state are due to crystal contacts. Thus, in that region the option to deactivate stage2 dihedral constraints will be used and fragments insertion will be performed.

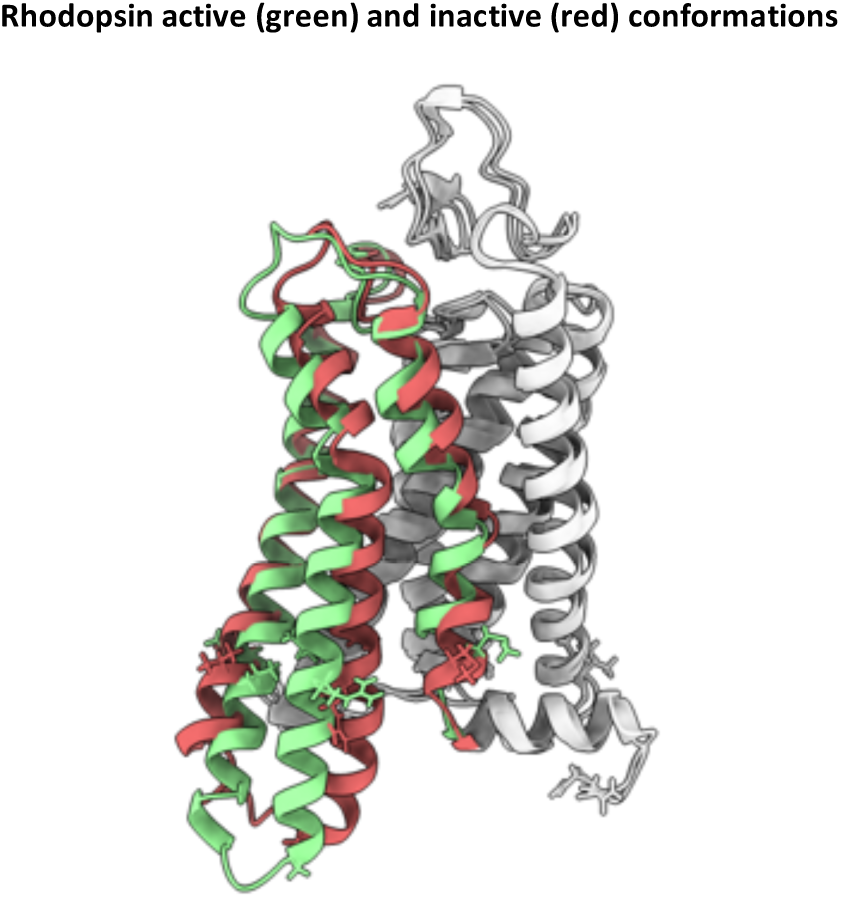

#### Preparation of the input files

The active (PDB ID 2×72) and inactive (PDB ID 1GZM) structures can be downloaded from ProteinDataBank. All the residues except range 3-326 of chain A are removed. In both the structures all the missing residues can be modeled by uploading sequence and template in the user-friendly portal https://swissmodel.expasy.org (click on “start modeling” followed by “User template: on the right).

Fragments can be collected through http://old.robetta.org. The website requires the protein fasta sequence to generate 3-mer and 9-mer fragments.

For membrane proteins, Rosetta requires a trans-membrane span topology file in which protein regions embedded in membrane are specified. There are two ways to generate a span file. 1) PDB file with membrane coordinates can be downloaded from the OPM webserver (http://www.opm.phar.umich.edu) and converted through the “mp_span_from_pdb” executable. 2) trans-membrane residues can be predicted with the OCTOPUS webserver at https://octopus.cbr.su.se and the file converted to the span format with the “octopus2span.pl” script located in the “scripts” folder.

Ten DEER distance distributions involving TM5-TM6-TM7 helices have been predicted with the DEER Spin-Pair Distributor in the CHARMM-GUI webserver at https://charmm-gui.org. The median value has been used in the restraints file.

All the following input files have been placed inside the “input_files” folder. “2×72.pdb” and “1gzm.pdb” for input active and target inactive structures, respectively. “aat000_03_05.200_v1_3.txt” and “aat000_09_05.200_v1_3.txt” for 3-mer and 9-mer fragments, respectively. The fasta sequence file “rhodopsin.fasta”. The span file “rhodopsin.span”. The restraints file “1gzm_simulated.cst”. A customized stage2_membrane scoring function “stage2_membrane.wts”.

#### RosettaScripts XML file

RosettaScripts XML file used for modeling has been deposited in the “scripts” folder and is shown below. The “modify_segments” option is used to include residue 306 in the segment 9 that otherwise would end at residue 305. Because residue 306 is restrained, its inclusion in a segment makes sure that such restraints will be fully exploited in stage 1. The “stage2_residues_no_dihedral_csts” is used to remove dihedral constraints of regions selected through “residue_selectors”. “stage1_multi_sse_freq” defines the frequency of performing multiple randomly chosen SSEs movements.

Besides ConfChangeMover, a couple of metrics are calculated: 1) the RMSD from the native structure of TM5-TM6-TM7 (_H suffix) and 2) the RMSD of the while modeled region (_HL suffix).

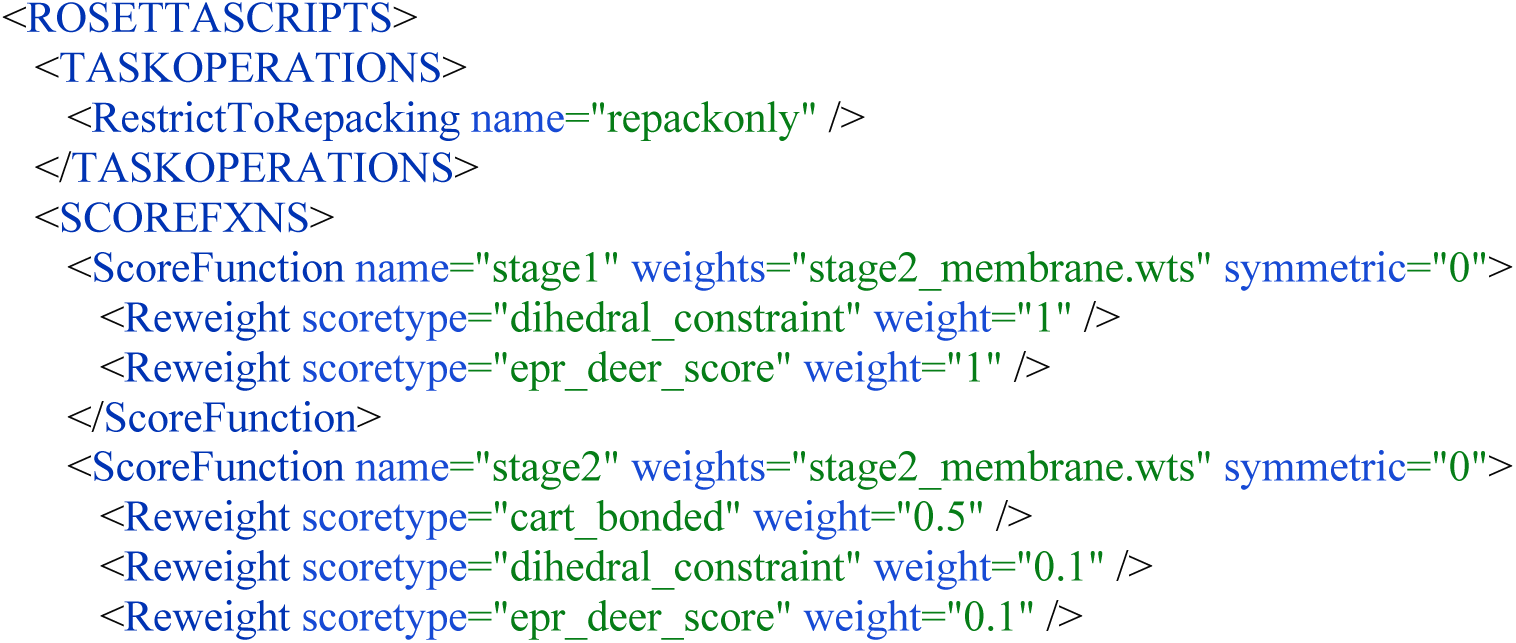
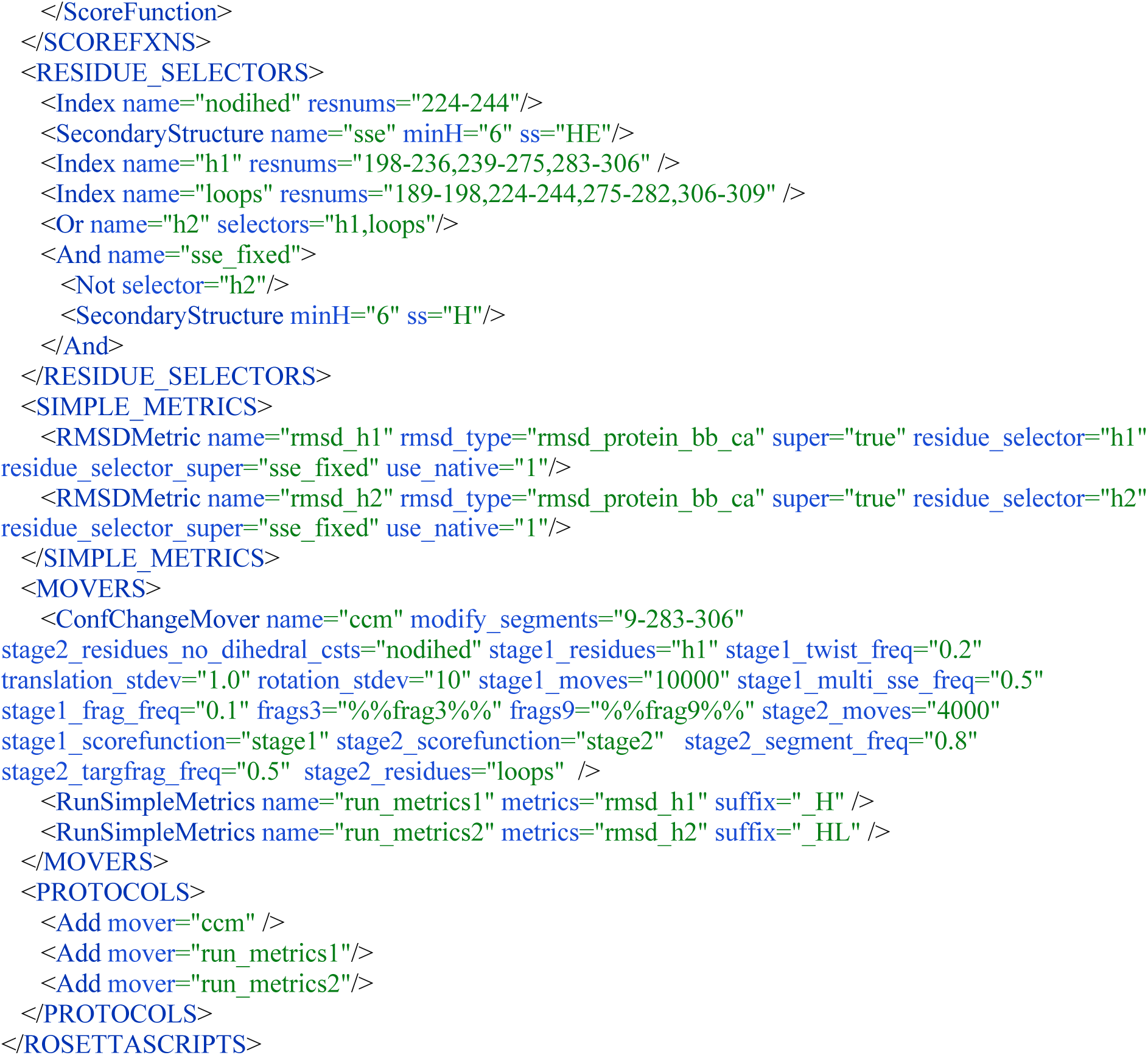

#### Running the command to perform the modeling

First, copy the scoring function *stage2_membrane*.*wts* in the “working_directory” folder. By executing the command below from “working_directory”, 5 models are generated in approximatively 45 minutes on a single core. User may need to change the executable depending on the OS used, MAC/linux/Windows. Rosetta executables are located in “path/to/Rosetta/main/source/bin”.

rosetta_scripts.linuxgccrelease -in:file:s ../input_files/2×72.pdb -in:file:native ../input_files/1gzm.pdb -in:file:spanfile ../input_files/rhodopsin.span -parser:script_vars frag3=../input_files/aat000_03_05.200_v1_3.txt frag9=../input_files/aat000_09_05.200_v1_3.txt -epr_deer:input_files ../input_files/1gzm_simulated.cst -parser:protocol ../scripts/ccm.xml -out:nstruct 5 - out:path:pdb ./ -mute core.optimization.Minimizer core.scoring.MembranePotential core.pose.util

#### Analyzing results in output

Depending on the number of models generated user can find the corresponding number of PDB files and a score file inside the “working_directory” folder. The score file contains the RMSD metrics for each model. Pre-generated output files can be found in the “output_files” folder.

**Figure S1.**
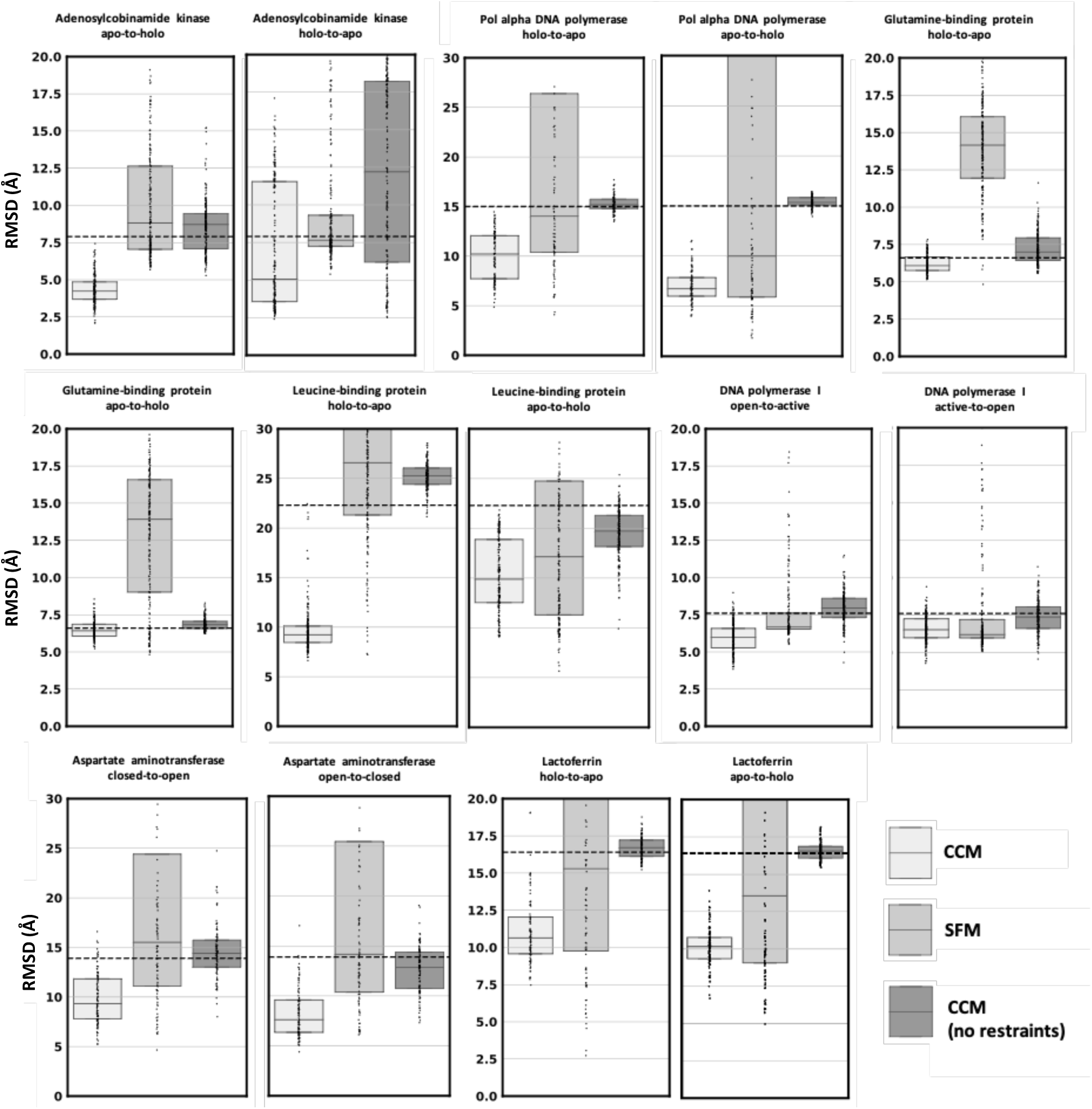
CCM performance in modeling conformational changes of SSEs and loops using simulated Cα-Cα distance restraints. Dots represent the real distribution of RMSD values. The RMSD between the two PDB conformations is represented as a dashed line.

**Figure S2.**
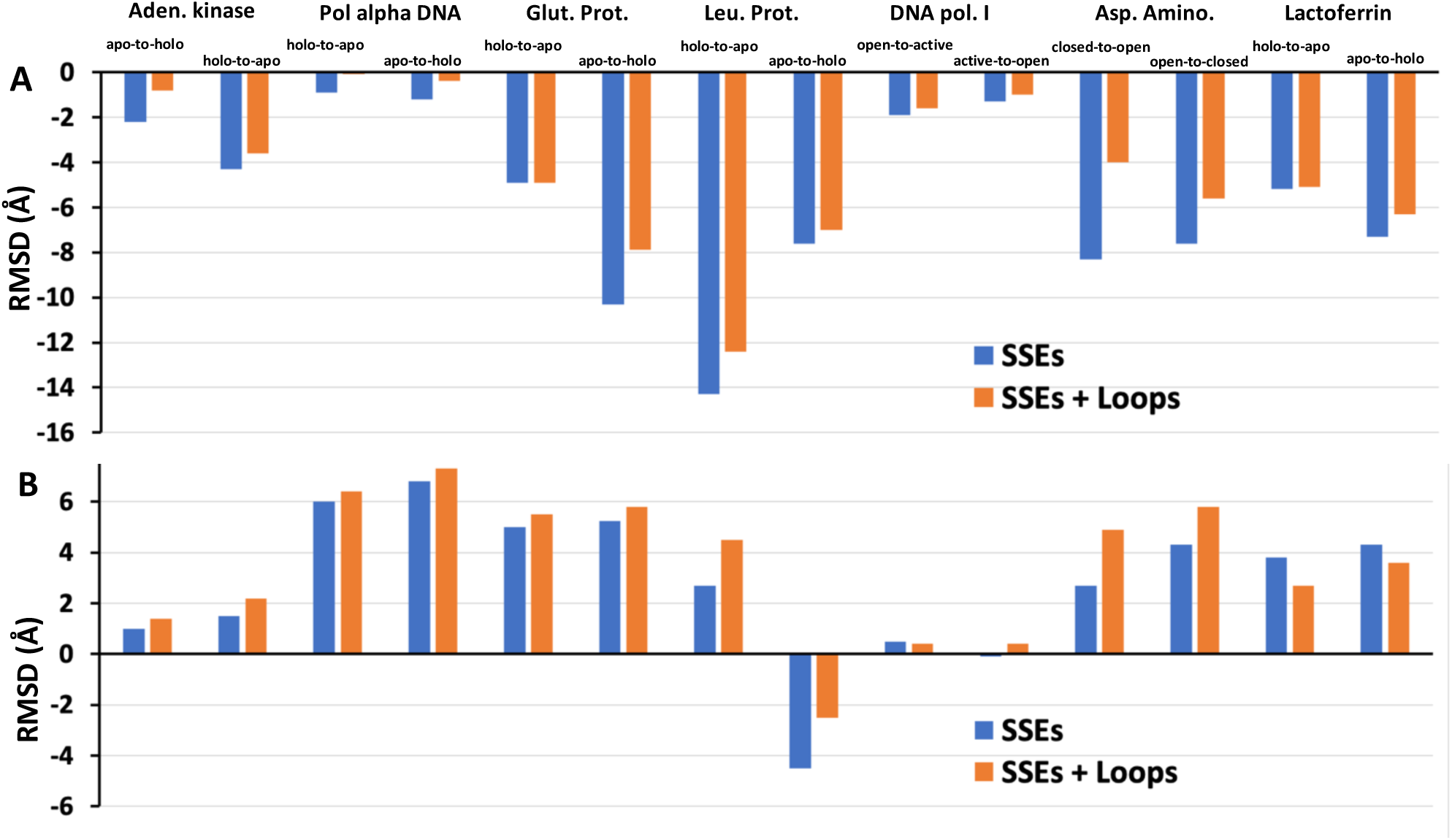
Change in RMSD of SSEs and loops. The RMSD difference was measured between median value from the target of each conformational ensemble and the value between experimental conformations. A) CCM. B) SFM.

**Figure S3.**
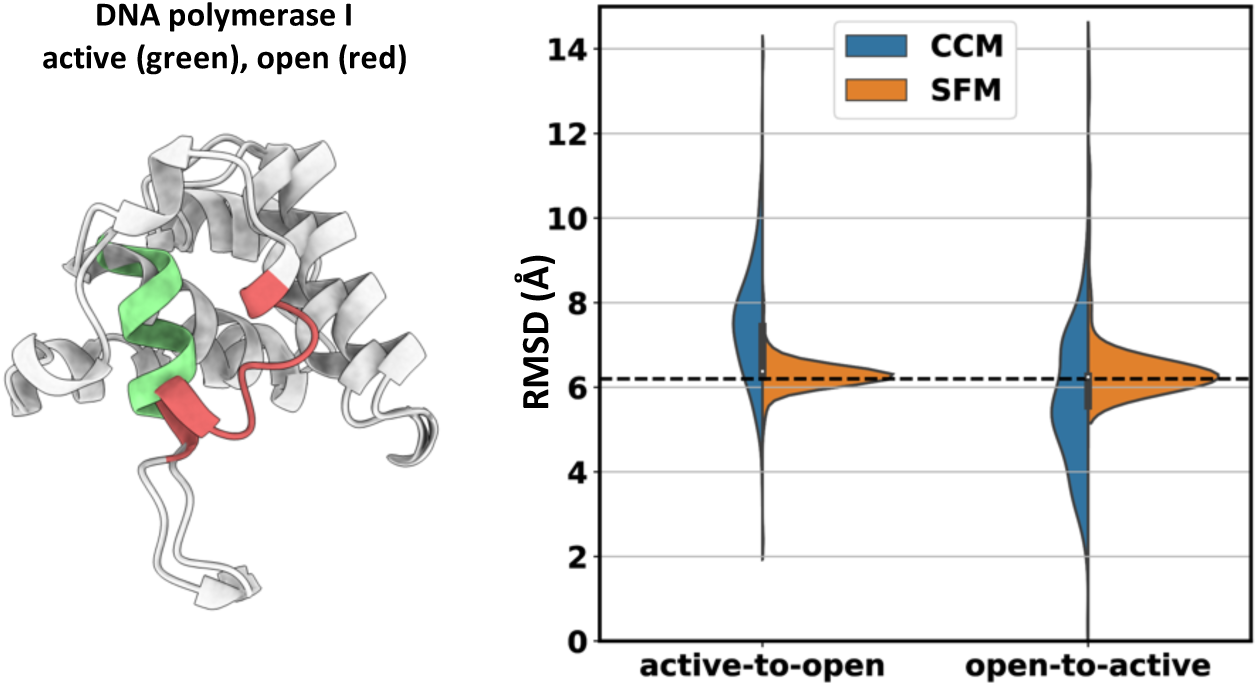
Accuracy in modeling a small helix with different folding states. Folding states are shown with colored cartoons. RMSD between native states is shown as a dashed line.

**Figure S4.**
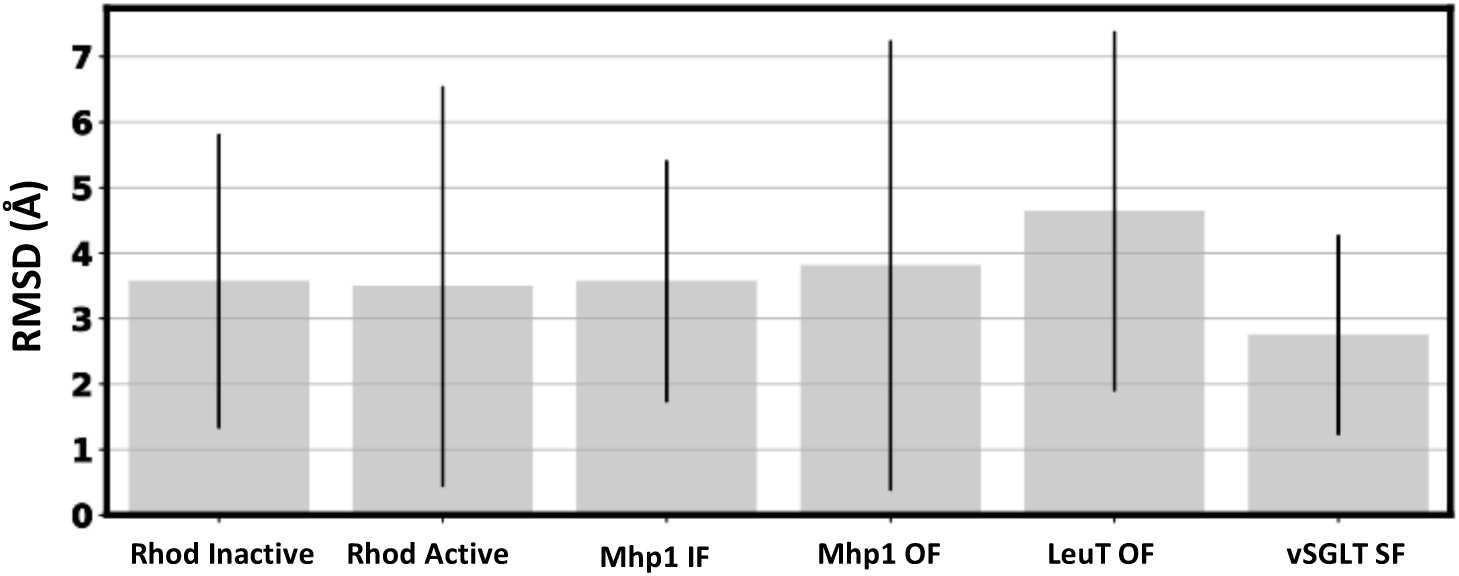
Accuracy of experimental DEER distances datasets. Bars indicate average RMSD between experimental and simulated DEER distances. Black lines represent standard deviations.

**Figure S5.**
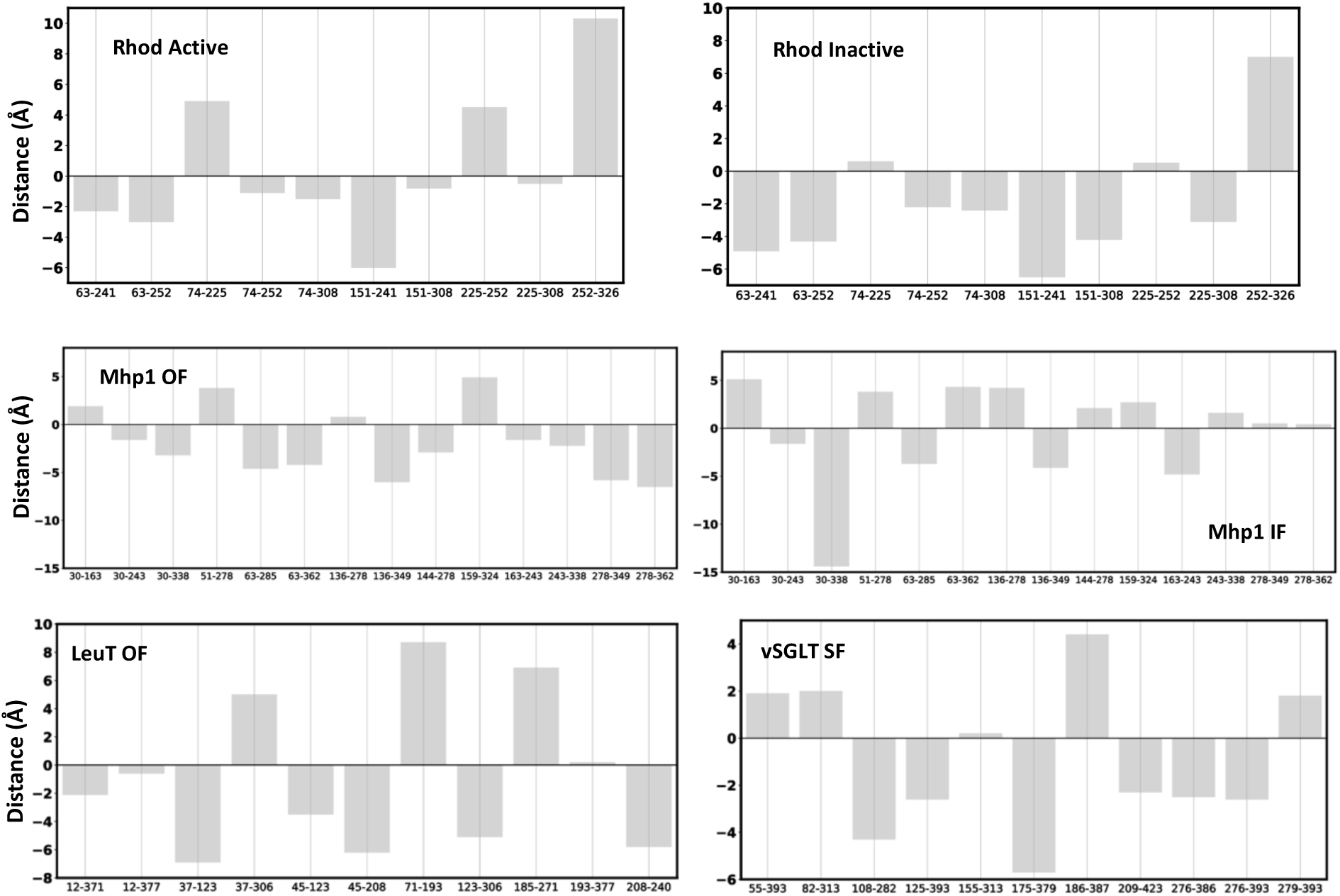
Accuracy of each experimental DEER distance restraint. The difference between experimental DEER distance and the corresponding simulated value was measured for each restrained residue pair.

**Figure S6.**
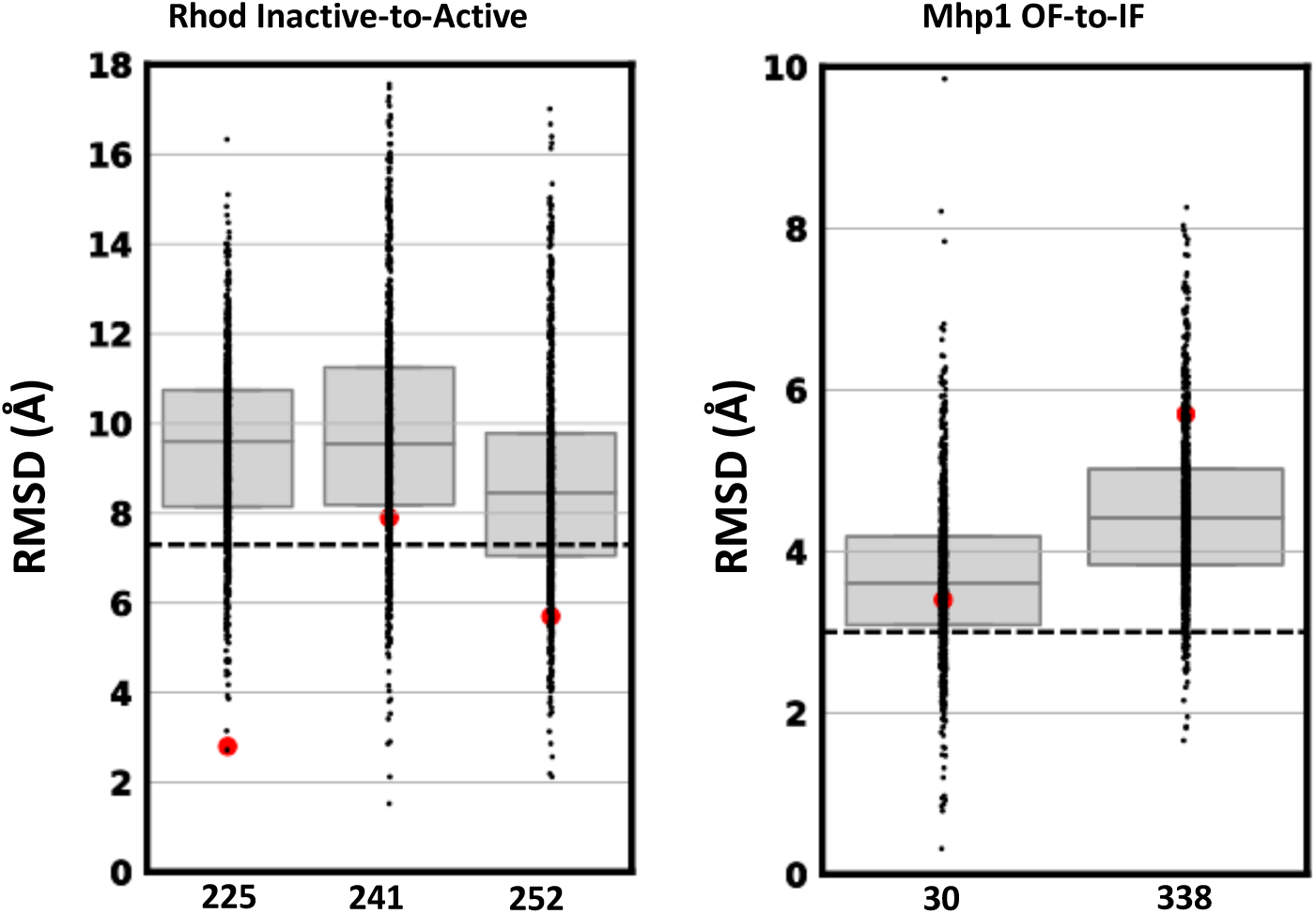
RMSD of residues involved in the most inaccurate experimental restraints of Rhodopsin and Mhp1 CCM (experimental median) ensembles. Black dots represent real data values. RMSD of the residue in the native starting conformation is shown as red dot. The median value of the global RMSD distribution of models is shown as dashed line.

## References

1. Nakane T, Kotecha A, Sente A, McMullan G, Masiulis S, Brown PMGE, et al. Single-particle cryo-EM at atomic resolution. Nature. 2020;587: 152–156. doi:10.1038/s41586-020-2829-0

2. Yip KM, Fischer N, Paknia E, Chari A, Stark H. Atomic-resolution protein structure determination by cryo-EM. Nature. 2020;587: 157–161. doi:10.1038/s41586-020-2833-4

3. Shimada I, Ueda T, Kofuku Y, Eddy MT, Wüthrich K. GPCR drug discovery: Integrating solution NMR data with crystal and cryo-EM structures. Nature Reviews Drug Discovery. Nature Publishing Group; 2018. pp. 59–82. doi:10.1038/nrd.2018.180

4. Baek M, DiMaio F, Anishchenko I, Dauparas J, Ovchinnikov S, Lee GR, et al. Accurate prediction of protein structures and interactions using a three-track neural network. Science (80-). 2021;373: 871–876. doi:10.1126/science.abj8754

5. Jumper J, Evans R, Pritzel A, Green T, Figurnov M, Ronneberger O, et al. Highly accurate protein structure prediction with AlphaFold. Nature. 2021;596: 583–589. doi:10.1038/s41586-021-03819-2

6. Tunyasuvunakool K, Adler J, Wu Z, Green T, Zielinski M, Žídek A, et al. Highly accurate protein structure prediction for the human proteome. Nature. 2021;596: 590–596. doi:10.1038/s41586-021-03828-1

7. Baek M, Anishchenko I, Park H, Humphreys IR, Baker D. Protein oligomer modeling guided by predicted interchain contacts in CASP14. Proteins Struct Funct Bioinforma. 2021. doi:10.1002/prot.26197

8. Schaeffer RD, Kinch L, Kryshtafovych A, Grishin N V. Assessment of domain interactions in CASP14. Proteins Struct Funct Bioinforma. 2021 [cited 24 Sep 2021]. doi:10.1002/prot.26225

9. Humphreys IR, Pei J, Baek M, Krishnakumar A, Anishchenko I, Ovchinnikov S, et al. Computed structures of core eukaryotic protein complexes. Science (80-). 2021 [cited 17 Nov 2021]. doi:10.1126/science.abm4805

10. Green AG, Elhabashy H, Brock KP, Maddamsetti R, Kohlbacher O, Marks DS. Large-scale discovery of protein interactions at residue resolution using co-evolution calculated from genomic sequences. Nat Commun. 2021;12: 1–12. doi:10.1038/s41467-021-21636-z

11. AlQuraishi M. Machine learning in protein structure prediction. Current Opinion in Chemical Biology. Elsevier Current Trends; 2021. pp. 1–8. doi:10.1016/j.cbpa.2021.04.005

12. Maximova T, Moffatt R, Ma B, Nussinov R, Shehu A. Principles and Overview of Sampling Methods for Modeling Macromolecular Structure and Dynamics. PLoS Computational Biology. 2016. doi:10.1371/journal.pcbi.1004619

13. Bernardi RC, Melo MCR, Schulten K. Enhanced sampling techniques in molecular dynamics simulations of biological systems. Biochimica et Biophysica Acta - General Subjects. Elsevier; 2015. pp. 872–877. doi:10.1016/j.bbagen.2014.10.019

14. Dastvan R, Fischer AW, Mishra S, Meiler J, McHaourab HS. Protonation-dependent conformational dynamics of the multidrug transporter EmrE. Proc Natl Acad Sci U S A. 2016;113: 1220–1225. doi:10.1073/pnas.1520431113

15. Bryn Fenwick R, Van Den Bedem H, Fraser JS, Wright PE. Integrated description of protein dynamics from room-temperature X-ray crystallography and NMR. Proc Natl Acad Sci U S A. 2014;111. doi:10.1073/pnas.1323440111

16. Greenleaf WJ, Woodside MT, Block SM. High-resolution, single-molecule measurements of biomolecular motion. Annual Review of Biophysics and Biomolecular Structure. Annu Rev Biophys Biomol Struct; 2007. pp. 171–190. doi:10.1146/annurev.biophys.36.101106.101451

17. Bonomi M, Heller GT, Camilloni C, Vendruscolo M. Principles of protein structural ensemble determination. Current Opinion in Structural Biology. Elsevier Ltd; 2017. pp. 106–116. doi:10.1016/j.sbi.2016.12.004

18. Palamini M, Canciani A, Forneris F. Identifying and visualizing macromolecular flexibility in structural biology. Frontiers in Molecular Biosciences. Frontiers Media S.A.; 2016. doi:10.3389/fmolb.2016.00047

19. Leaver-Fay A, Tyka M, Lewis SM, Lange OF, Thompson J, Jacak R, et al. Rosetta3: An object-oriented software suite for the simulation and design of macromolecules. Methods in Enzymology. Methods Enzymol; 2011. pp. 545–574. doi:10.1016/B978-0-12-381270-4.00019-6

20. Eswar N, Webb B, Marti-Renom MA, Madhusudhan MS, Eramian D, Shen M, et al. Comparative Protein Structure Modeling Using MODELLER. Current Protocols in Protein Science. Hoboken, NJ, USA: John Wiley & Sons, Inc.; 2007. pp. 2.9.1-2.9.31. doi:10.1002/0471140864.ps0209s50

21. Dominguez C, Boelens R, Bonvin Amjj. HADDOCK: A protein-protein docking approach based on biochemical or biophysical information. J Am Chem Soc. 2003;125: 1731–1737. doi:10.1021/ja026939x

22. Xia Y, Fischer AW, Teixeira P, Weiner B, Meiler J. Integrated Structural Biology for α-Helical Membrane Protein Structure Determination. Structure. 2018;26: 657-666.e2. doi:10.1016/j.str.2018.02.006

23. Leman JK, Weitzner BD, Lewis SM, Adolf-Bryfogle J, Alam N, Alford RF, et al. Macromolecular modeling and design in Rosetta: recent methods and frameworks. Nat Methods. 2020. doi:10.1038/s41592-020-0848-2

24. Heilmann N, Wolf M, Kozlowska M, Sedghamiz E, Setzler J, Brieg M, et al. Sampling of the conformational landscape of small proteins with Monte Carlo methods. Sci Rep. 2020;10: 1–13. doi:10.1038/s41598-020-75239-7

25. Park H, Ovchinnikov S, Kim DE, DiMaio F, Baker D. Protein homology model refinement by large-scale energy optimization. Proc Natl Acad Sci U S A. 2018;115: 3054–3059. doi:10.1073/pnas.1719115115

26. Song Y, Dimaio F, Wang RYR, Kim D, Miles C, Brunette T, et al. High-resolution comparative modeling with RosettaCM. Structure. 2013;21: 1735–1742. doi:10.1016/j.str.2013.08.005

27. Sfriso P, Duran-Frigola M, Mosca R, Emperador A, Aloy P, Orozco M. Residues Coevolution Guides the Systematic Identification of Alternative Functional Conformations in Proteins. Structure. 2016;24: 116–126. doi:10.1016/j.str.2015.10.025

28. Jeschke G. Characterization of protein conformational changes with sparse spin-label distance constraints. J Chem Theory Comput. 2012;8: 3854–3863. doi:10.1021/ct300113z

29. Zheng W, Brooks BR. Normal-modes-based prediction of protein conformational changes guided by distance constraints. Biophys J. 2005;88: 3109–3117. doi:10.1529/biophysj.104.058453

30. Kim DE, Chivian D, Baker D. Protein structure prediction and analysis using the Robetta server. Nucleic Acids Res. 2004;32. doi:10.1093/nar/gkh468

31. Thompson TB, Thomas MG, Escalante-Semerena JC, Rayment I. Three-dimensional structure of adenosylcobinamide kinase/adenosylcobinamide phosphate guanylyltransferase (CobU) complexed with GMP: Evidence for a substrate-induced transferase active site. Biochemistry. 1999;38: 12995–13005. doi:10.1021/bi990910x

32. Thompson TB, Thomas MG, Escalante-Semerena JC, Rayment I. Three-dimensional structure of adenosylcobinamide kinase/adenosylcobinamide phosphate guanylyltransferase from Salmonella typhimurium determined to 2.3 Å resolution. Biochemistry. 1998;37: 7686–7695. doi:10.1021/bi973178f

33. Franklin MC, Wang J, Steitz TA. Structure of the replicating complex of a pol α family DNA polymerase. Cell. 2001;105: 657–667. doi:10.1016/S0092-8674(01)00367-1

34. Sun YJ, Rose J, Wang BC, Hsiao CD. The structure of glutamine-binding protein complexed with glutamine at 1.94 Å resolution: Comparisons with other amino acid binding proteins. J Mol Biol. 1998;278: 219–229. doi:10.1006/jmbi.1998.1675

35. Hsiao CD, Sun YJ, Rose J, Wang BC. The crystal structure of glutamine-binding protein from Escherichia coli. J Mol Biol. 1996;262: 225–242. doi:10.1006/jmbi.1996.0509

36. Magnusson U, Salopek-Sondi B, Luck LA, Mowbray SL. X-ray Structures of the Leucine-binding Protein Illustrate Conformational Changes and the Basis of Ligand Specificity. J Biol Chem. 2004;279: 8747–8752. doi:10.1074/jbc.M311890200

37. Li Y, Korolev S, Waksman G. Crystal structures of open and closed forms of binary and ternary complexes of the large fragment of Thermus aquaticus DNA polymerase I: Structural basis for nucleotide incorporation. EMBO J. 1998;17: 7514–7525. doi:10.1093/emboj/17.24.7514

38. McPhalen CA, Vincent MG, Picot D, Jansonius JN, Lesk AM, Chothia C. Domain closure in mitochondrial aspartate aminotransferase. J Mol Biol. 1992;227: 197–213. doi:10.1016/0022-2836(92)90691-C

39. McPhalen CA, Vincent MG, Jansonius JN. X-ray structure refinement and comparison of three forms of mitochondrial aspartate aminotransferase. J Mol Biol. 1992;225: 495–517. doi:10.1016/0022-2836(92)90935-D

40. Norris GE, Anderson BF, Baker EN. Molecular replacement solution of the structure of apolactoferrin, a protein displaying large-scale conformational change. Acta Crystallogr Sect B. 1991;47: 998–1004. doi:10.1107/S0108768191008418

41. Haridas M, Anderson BF, Baker EN. Structure of human diferrric lactoferrin refined at 2.2 Angstrom resolution. Acta Crystallogr - Sect D Biol Crystallogr. 1995;51: 629–646. doi:10.1107/S0907444994013521

42. Yamashita A, Singh SK, Kawate T, Jin Y, Gouaux E. Crystal structure of a bacterial homologue of Na+/Cl --dependent neurotransmitter transporters. Nature. 2005;437: 215–223. doi:10.1038/nature03978

43. Krishnamurthy H, Gouaux E. X-ray structures of LeuT in substrate-free outward-open and apo inward-open states. Nature. 2012;481: 469–474. doi:10.1038/nature10737

44. Kazmier K, Sharma S, Quick M, Islam SM, Roux B, Weinstein H, et al. Conformational dynamics of ligand-dependent alternating access in LeuT. Nat Struct Mol Biol. 2014;21: 472–479. doi:10.1038/nsmb.2816

45. Shimamura T, Weyand S, Beckstein O, Rutherford NG, Hadden JM, Sharpies D, et al. Molecular basis of alternating access membrane transport by the sodium-hydantoin transporter Mhp1. Science (80-). 2010;328: 470–473. doi:10.1126/science.1186303

46. Weyand S, Shimamura T, Yajima S, Suzuki SNI, Mirza O, Krusong K, et al. Structure and molecular mechanism of a nucleobase-cation-symport-1 family transporter. Science (80-). 2008;322: 709–713. doi:10.1126/science.1164440

47. Wahlgren WY, Dunevall E, North RA, Paz A, Scalise M, Bisignano P, et al. Substrate-bound outward-open structure of a Na+-coupled sialic acid symporter reveals a new Na+ site. Nat Commun. 2018;9: 1–14. doi:10.1038/s41467-018-04045-7

48. Watanabe A, Choe S, Chaptal V, Rosenberg JM, Wright EM, Grabe M, et al. The mechanism of sodium and substrate release from the binding pocket of vSGLT. Nature. 2010;468: 988–991. doi:10.1038/nature09580

49. Paz A, Claxton DP, Kumar JP, Kazmier K, Bisignano P, Sharma S, et al. Conformational transitions of the sodium-dependent sugar transporter, vSGLT. Proc Natl Acad Sci U S A. 2018;115: E2742–E2751. doi:10.1073/pnas.1718451115

50. Standfuss J, Edwards PC, D’Antona A, Fransen M, Xie G, Oprian DD, et al. The structural basis of agonist-induced activation in constitutively active rhodopsin. Nature. 2011;471: 656–660. doi:10.1038/nature09795

51. Li J, Edwards PC, Burghammer M, Villa C, Schertler GFX. Structure of bovine rhodopsin in a trigonal crystal form. J Mol Biol. 2004;343: 1409–1438. doi:10.1016/j.jmb.2004.08.090

52. Islam SM, Roux B. Simulating the distance distribution between spin-labels attached to proteins. J Phys Chem B. 2015;119: 3901–3911. doi:10.1021/jp510745d

53. Islam SM, Stein RA, McHaourab HS, Roux B. Structural refinement from restrained-ensemble simulations based on EPR/DEER data: Application to T4 lysozyme. J Phys Chem B. 2013;117: 4740–4754. doi:10.1021/jp311723a

54. Qi Y, Lee J, Cheng X, Shen R, Islam SM, Roux B, et al. CHARMM-GUI DEER facilitator for spin-pair distance distribution calculations and preparation of restrained-ensemble molecular dynamics simulations. J Comput Chem. 2020;41: 415–420. doi:10.1002/jcc.26032

55. Jo S, Kim T, Iyer VG, Im W. CHARMM-GUI: A web-based graphical user interface for CHARMM. J Comput Chem. 2008;29: 1859–1865. doi:10.1002/jcc.20945

56. Claxton DP, Quick M, Shi L, De Carvalho FD, Weinstein H, Javitch JA, et al. Ion/substrate-dependent conformational dynamics of a bacterial homolog of neurotransmitter:sodium symporters. Nat Struct Mol Biol. 2010;17: 822–829. doi:10.1038/nsmb.1854

57. Kazmier K, Sharma S, Islam SM, Roux B, Mchaourab HS, Wright EM. Conformational cycle and ion-coupling mechanism of the Na+/hydantoin transporter Mhp1. Proc Natl Acad Sci U S A. 2014;111: 14752–14757. doi:10.1073/pnas.1410431111

58. del Alamo D, Tessmer MH, Stein RA, Feix JB, Mchaourab HS, Meiler J. Rapid Simulation of Unprocessed DEER Decay Data for Protein Fold Prediction. Biophys J. 2020;118: 366–375. doi:10.1016/j.bpj.2019.12.011

59. Viklund H, Elofsson A. OCTOPUS: Improving topology prediction by two-track ANN-based preference scores and an extended topological grammar. Bioinformatics. 2008;24: 1662–1668. doi:10.1093/bioinformatics/btn221

60. Yarov-Yarovoy V, Schonbrun J, Baker D. Multipass membrane protein structure prediction using Rosetta. Proteins Struct Funct Genet. 2006;62: 1010–1025. doi:10.1002/prot.20817

61. Fleishman SJ, Leaver-Fay A, Corn JE, Strauch EM, Khare SD, Koga N, et al. Rosettascripts: A scripting language interface to the Rosetta Macromolecular modeling suite. PLoS One. 2011;6: e20161. doi:10.1371/journal.pone.0020161

62. Kabsch W, Sander C. Dictionary of protein secondary structure: Pattern recognition of hydrogen-bonded and geometrical features. Biopolymers. 1983;22: 2577–2637. doi:10.1002/bip.360221211

63. Tyka MD, Jung K, Baker D. Efficient sampling of protein conformational space using fast loop building and batch minimization on highly parallel computers. J Comput Chem. 2012;33: 2483–2491. doi:10.1002/jcc.23069

64. Ovchinnikov S, Kinch L, Park H, Liao Y, Pei J, Kim DE, et al. Large-scale determination of previously unsolved protein structures using evolutionary information. Elife. 2015;4: 1–25. doi:10.7554/eLife.09248

65. Woetzel N, Karakas M, Staritzbichler R, Müller R, Weiner BE, Meiler J. BCL::Score-Knowledge Based Energy Potentials for Ranking Protein Models Represented by Idealized Secondary Structure Elements. PLoS One. 2012;7: e49242. doi:10.1371/journal.pone.0049242

66. Rohl CA, Strauss CEM, Chivian D, Baker D. Modeling Structurally Variable Regions in Homologous Proteins with Rosetta. Proteins Struct Funct Genet. 2004;55: 656–677. doi:10.1002/prot.10629

67. Conway P, Tyka MD, DiMaio F, Konerding DE, Baker D. Relaxation of backbone bond geometry improves protein energy landscape modeling. Protein Sci. 2014;23: 47–55. doi:10.1002/pro.2389

68. Byrd RH, Lu P, Nocedal J, Zhu C. A Limited Memory Algorithm for Bound Constrained Optimization. SIAM J Sci Comput. 1995;16: 1190–1208. doi:10.1137/0916069

69. Bystroff C, Baker D. Prediction of local structure in proteins using a library of sequence-structure motifs. J Mol Biol. 1998;281: 565–577. doi:10.1006/jmbi.1998.1943

70. Simons KT, Kooperberg C, Huang E, Baker D. Assembly of protein tertiary structures from fragments with similar local sequences using simulated annealing and Bayesian scoring functions. J Mol Biol. 1997;268: 209–225. doi:10.1006/jmbi.1997.0959

71. Park H, Lee GR, Kim DE, Anishchenko I, Cong Q, Baker D. High-accuracy refinement using Rosetta in CASP13. Proteins Struct Funct Bioinforma. 2019;87: 1276–1282. doi:10.1002/prot.25784

72. Hiranuma N, Park H, Baek M, Anishchenko I, Dauparas J, Baker D. Improved protein structure refinement guided by deep learning based accuracy estimation. Nat Commun. 2021;12: 1–11. doi:10.1038/s41467-021-21511-x

73. Kuenze G, Bonneau R, Leman JK, Meiler J. Integrative Protein Modeling in RosettaNMR from Sparse Paramagnetic Restraints. Structure. 2019;27: 1721-1734.e5. doi:10.1016/j.str.2019.08.012

74. Evans EGB, Morgan JLW, DiMaio F, Zagotta WN, Stoll S. Allosteric conformational change of a cyclic nucleotide-gated ion channel revealed by DEER spectroscopy. Proc Natl Acad Sci U S A. 2020;117: 10839–10847. doi:10.1073/PNAS.1916375117

75. Pilla KB, Leman JK, Otting G, Huber T. Capturing conformational states in proteins using sparse paramagnetic NMR data. PLoS One. 2015;10. doi:10.1371/journal.pone.0127053

76. del Alamo D, Fischer AW, Moretti R, Alexander NS, Mendenhall J, Hyman NJ, et al. Efficient Sampling of Protein Loop Regions Using Conformational Hashing Complemented with Random Coordinate Descent. J Chem Theory Comput. 2021;17: 560–570. doi:10.1021/acs.jctc.0c00836

77. Stein A, Kortemme T. Improvements to Robotics-Inspired Conformational Sampling in Rosetta. PLoS One. 2013;8: e63090. doi:10.1371/journal.pone.0063090

78. Mandell DJ, Coutsias EA, Kortemme T. Sub-angstrom accuracy in protein loop reconstruction by robotics-inspired conformational sampling. Nat Methods. 2009;6: 551–552. doi:10.1038/nmeth0809-551

79. Canutescu AA, Dunbrack RL. Cyclic coordinate descent: A robotics algorithm for protein loop closure. Protein Sci. 2003;12: 963–972. doi:10.1110/ps.0242703

80. Hays JM, Kieber MK, Li JZ, Han JI, Columbus L, Kasson PM. Refinement of Highly Flexible Protein Structures using Simulation-Guided Spectroscopy. Angew Chemie - Int Ed. 2018;57: 17110–17114. doi:10.1002/anie.201810462

81. Marzolf DR, Seffernick JT, Lindert S. Protein Structure Prediction from NMR Hydrogen-Deuterium Exchange Data. J Chem Theory Comput. 2021;17: 2619–2629. doi:10.1021/acs.jctc.1c00077

82. Biehn SE, Limpikirati P, Vachet RW, Lindert S. Utilization of Hydrophobic Microenvironment Sensitivity in Diethylpyrocarbonate Labeling for Protein Structure Prediction. Anal Chem. 2021;93: 8188–8195. doi:10.1021/acs.analchem.1c00395

83. Aprahamian ML, Chea EE, Jones LM, Lindert S. Rosetta Protein Structure Prediction from Hydroxyl Radical Protein Footprinting Mass Spectrometry Data. Anal Chem. 2018;90: 7721–7729. doi:10.1021/acs.analchem.8b01624

84. Feng J, Shukla D. Characterizing Conformational Dynamics of Proteins Using Evolutionary Couplings. J Phys Chem B. 2018;122: 1017–1025. doi:10.1021/acs.jpcb.7b07529

